# Balancing control: a Bayesian interpretation of habitual and goal-directed behavior

**DOI:** 10.1101/836106

**Authors:** Sarah Schwöbel, Dimitrije Markovic, Michael N. Smolka, Stefan J. Kiebel

## Abstract

In everyday life, our behavior varies on a continuum from automatic and habitual to deliberate and goal-directed. Recent evidence suggests that habit formation and relearning of habits operate in a context-dependent manner: Habit formation is promoted when actions are performed in a specific context, while breaking off habits is facilitated after a context change. It is an open question how one can computationally model the brain’s balancing between context-specific habits and goal-directed actions. Here, we propose a hierarchical Bayesian approach for control of a partially observable Markov decision process that enables conjoint learning of habits and reward structure in a context-specific manner. In this model, habit learning corresponds to an updating of priors over policies and interacts with the learning of the outcome contingencies. Importantly, the model is solely built on probabilistic inference, which effectively provides a simple explanation of how the brain may balance contributions of habitual and goal-directed control. We illustrated the resulting behavior using agent-based simulated experiments, where we replicated several findings of devaluation, extinction, and renewal experiments, as well as the so-called two-step task which is typically used with human participants. In addition, we show how a single parameter, the habitual tendency, can explain individual differences in habit learning and the balancing between habitual and goal-directed control. Finally, we discuss the link of the proposed model to other habit learning models and implications for understanding specific phenomena in substance use disorder.

## 1. Introduction

In both psychology and neuroscience, theories postulate that behavioral control can vary along a dimension with habitual, automatic actions on one end, and goal-directed, controlled actions on the other (Wood and Rünger, 2016). In the context of operant conditioning, habits have been described as retrospective and have been found to implement an automatic tendency to repeat actions which have been rewarded in the past (Dickinson et al., 1983; Graybiel, 2008). Habitual action selection is typically fast but insensitive to outcomes and only slowly adapts to a changing environment (Seger and Spiering, 2011). In contrast, goal-directed action selection is prospective and implements planning based on a representation of action-outcome contingencies (Dickinson and Balleine, 1994; Dolan and Dayan, 2013). Consequently, goal-directed action selection adapts rather rapidly to a changing environment, but under a penalty of costly and slow computations.

Habit learning can be viewed as a transition from goal-directed to habitual behavior while a subject learns about its environment (Graybiel, 2008): In a novel environment or context, goal-directed actions will first allow the organism to learn about its structure and rewards and, later, to integrate this information to reliably reach a goal. With time, certain behaviors will be reinforced, while others will not. Subsequently, habits are formed to enable faster and computationally less costly selection of behavior which has been successful in the past. Given enough training, behavior is thought to be dominated by stimulus-driven habits, see e.g. (Dickinson, 1985; Seger and Spiering, 2011) for experimentally derived criteria of habit learning. In particular, two influential criteria are the insensitivity to contingency degradation where action-outcome associations are changed, and the insensitivity to reinforcer devaluation, where the outcome is made undesirable (Yin and Knowlton, 2006). Here, an established habit seems to make it difficult for an organism to change the previously reinforced habitual choice and adapt behavior to the altered conditions in its environment. Additionally, the strength of the habit and resulting insensitivity to changes has been found to critically depend on the duration and reward schedule of the training phase (Yin and Knowlton, 2006).

Importantly, habit learning as well as changing existing habits is strongly associated with the consistency of the environment while actions are performed (Wood and Rünger, 2016). When a specific behavior is executed in a stable context, habits are learned faster, and adjustment of behavioral patterns after changes in context is impeded (Lally et al., 2010). Conversely, learning of habits is slower and adjustment to changes is facilitated in a changing environment or inconsistent contexts. For example, it has been shown that learning of habits is improved when actions are mostly performed in the same context, e.g. after breakfast (Lally et al., 2010; Danner et al., 2008; Neal et al., 2012); while the unlearning of habits is improved after a context change, e.g. after a move to a different city (Verplanken and Roy, 2016).

In addition, habit learning trajectories strongly vary between individuals (Dolan and Dayan, 2013; Lally et al., 2010). Recent substance use disorder (SUD) studies show differences, between patients and controls, in learning and in the reliance on the so-called habit system, which lead to individual habitual responding biases (Ersche et al., 2016; Lim et al., 2019; Heinz et al., 2019). Still, it is an open question whether these different habit learning trajectories in individuals with SUD are due to individual factors or caused by the substance use itself (Nebe et al., 2018).

While there are findings that there are two hypothesized systems, the habitual and goal-directed system, and how they map onto brain structures (Dolan and Dayan, 2013; Yin and Knowlton, 2006; Everitt and Robbins, 2005), it is not clear if such a dichotomy is required for the computational description of these processes and for a mechanistic understanding of how habitual and goal-directed control are balanced, e.g. (Goschke, 2014). It has been argued that goal-directed and habitual behavior can be equated to model-based and model-free reinforcement learning (Daw et al., 2005, 2011; Dolan and Dayan, 2013), where arbitration may be based on the respective (un)certainties of the two hypothesized controllers. However, experimental evidence indicates that model-free reinforcement learning does not capture all experimentally established properties of habit formation (Friedel et al., 2014; Gillan et al., 2015). Rather, an alternative proposal is centered on the idea that habits, as stimulus-response associations, may arise from repetition alone and are learned via a value-free mechanism (Miller et al., 2019). Another emerging research direction, built on both experimental and computational studies, is to consider habits as chunked action sequences, which may be modelled in a hierarchical fashion (Smith and Graybiel, 2016; Graybiel and Grafton, 2015; Dezfouli and Balleine, 2012, 2013; Graybiel and Grafton, 2015).

Here, we want to build on these previous lines of work and present a hierarchical Bayesian ‘prior-based control’ habit-learning model. We build our modeling approach upon four assumptions: (i) Habits are learned based on repetition of past behavior. (ii) Habits are learned as chunked action sequences. (iii) Habitual and goal-directed control contributions are weighted by their respective (un)certainties. (iv) Habits and action-outcome contingencies are learned in a hierarchical, context-dependent manner.

To formally implement these assumptions, we build our model based on the concept of planning as inference (Attias, 2003; Botvinick and Toussaint, 2012), where action selection is regarded as an inference problem. The inference will be treated using approximate methods similar to the active inference framework (Friston et al., 2015), which we use to simulate agent behavior. We propose to view balancing of behavioral control in a Bayesian way: Behavior is sampled from a posterior which, according to Bayes’ rule, is proportional to a prior times a likelihood. We interpret a sharp or pronounced prior as the basis of a habit, see also (Friston et al., 2016), where the habitual contribution for a specific action or policy (sequence of actions) is higher the more this action, or sequence of actions, has been selected in the past (assumptions i and ii). The goal-directed control contribution of an action or policy is encoded in the likelihood, where explicit forward planning yields the expected reward. This explicit forward planning is based on a partially observable Markov decision process where policies enable navigation through a state space, and where outcome contingencies are learned. As a result, the interpretation of how control is balanced is rather simple: Goal-directed and habitual control contributions are multiplied using Bayes’ rule, yielding a natural weighting based on the respective certainties (assumption iii). Importantly, the prior, the outcome rules, and consequently the generative model underlying the partially observable Markov decision process, are learned in a context-specific manner (assumption iv). As a result, habits and outcome contingencies are acquired for each context and can be retrieved when re-encountering a known context. We use this hierarchical model to explain the transition dynamics from goal-directed to habitual behavior when learning habits, and adaptation of behavior to context changes.

We show using simulations that the prior-based control model is in principle able to capture basic properties of classical habit learning experiments: Insensitivity to changes in action-outcome contingencies and reinforcer devaluation, and the increase of this effect with longer training duration. We introduce a trait-like free parameter of the model, the habitual tendency, which modulates an individual’s habit learning speed. We also show that stochastic environments which are akin to interval reward schedules result in an over-reliance on habitual control. Furthermore, we illustrate that context-specific habits enable rapid adaptation after a switch to another but already known context, as typically found in ABA renewal experiments Bouton and Bolles (1979); Nakajima et al. (2000); Bouton et al. (2011). Finally, we show that the model replicates behavioral patterns as experimentally found in the sequential two-step task (Daw et al., 2011). Lastly, we will discuss the interpretations and implications of our model and how the proposal of habits modelled as priors over action sequences lets us reinterpret the assumed dichotomy of the habitual and goal-directed system.

## 2. Methods

Before we go into the details of the formal model, we sketch the key idea here. Under planning as inference, planning and action selection are understood as a Bayesian inference process

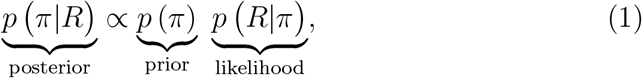

where actions, or policies *π* (sequences of actions) are part of the generative model and which action to take is a question that needs to be inferred. An agent evaluates the likelihood *p*(*R*|*π*) of obtaining rewards *R* under a certain policy *π*, which corresponds to goal-directed forward planning. The agent uses Bayes’ rule to evaluate which policy to follow given that it wants to achieve a reward in the posterior over actions *p*(*π*|*R*), from which an action is then sampled. The prior over policies *p*(*π*), is often overlooked or set to be flat, as it does not contribute to the goal-reaching behavior.

In this work, we posit that habitual control contributions correspond to the relative values in the prior over policies, while goal-directed contributions stem from the likelihood, both of which are automatically weighted by a simple multiplication to achieve a global control signal in the posterior. In this framework, a pronounced prior which is close to one for a specific policy and close to zero for all other policies corresponds to a strong habitual control contribution for this policy. Conversely, a flat prior, where all policies have the same prior value, corresponds to minimal habitual control contributions, as in this case the posterior is dominated by the likelihood. We term this kind of model ‘prior-based control model’. Furthermore, this approach allows to use Bayesian updating for the learning of the habitual control contributions in the prior, which yield a repetition-based learning rule. This simple formalism efficiently implements assumptions (i), (ii), and (iii). Lastly, we extend this formalism by a context *c*

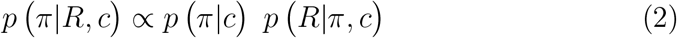

to include context-dependent learning of the prior as well as action-outcome contingencies in the likelihood (assumption iv). Below we will first introduce the generative process of the environment, and then introduce the full generative model which includes states an agent navigates through that link actions and rewards. We finally invert the full generative model using variational methods to obtain the agent’s beliefs and how they are updated.

### 2.1. The generative process

In this subsection, we describe the structure of the task environment. Our description rests on a hierarchical partially observable Markov decision process (POMDP), which is defined by the tuple 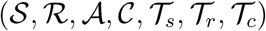, where

- 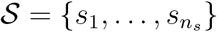 is a set of states
- 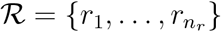 is a set of rewards
- 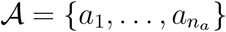 is a set of actions
- 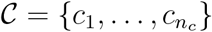 is a set of contexts
- 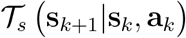 is a set of action-dependent state transition rules
- 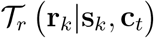 is a set of context-dependent reward generation rules
- 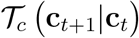 is a set of context transition rules.

Here, we deviate slightly from the typical notation used in the reinforcement learning literature on POMDPs, e.g. (Littman, 2009; Sutton and Barto, 1998), and instead adhere to the notation used in the active inference literature (Friston et al., 2015, 2016), where state transitions and reward generation are treated using separate transition rules. We partition the time evolution of the environment into *N_e_* episodes of length *K* (Hommel et al., 2001; Zacks et al., 2007; Butz, 2016). In the *t*-th episode, the environment is in context 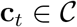. In this episode, the first time step is *k* = 1. The environment starts out in its starting state 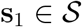. Depending on the state and the current context, the environment distributes a reward 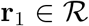 according to the generation rule 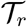, which essentially encodes the contingency tables for each context. Note that a no-reward is also part of the set of rewards 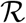. This way, the environment is set up to have a context-dependent reward distribution rule, which may also change, when the environment transitions to a new context. Using these transitions, we will be able to implement the training and extinction phases of a typical habit learning environment as latent contexts in the Markov decision process.

A subject or agent, which is interacting with this environment, observes the reward and state of the environment, and chooses an action **a**_1_. This marks the end of the first time step *k* = 1 of the *t*-th episode. This process for a single time step is also shown in the left part of Figure 1.

**Figure 1:**
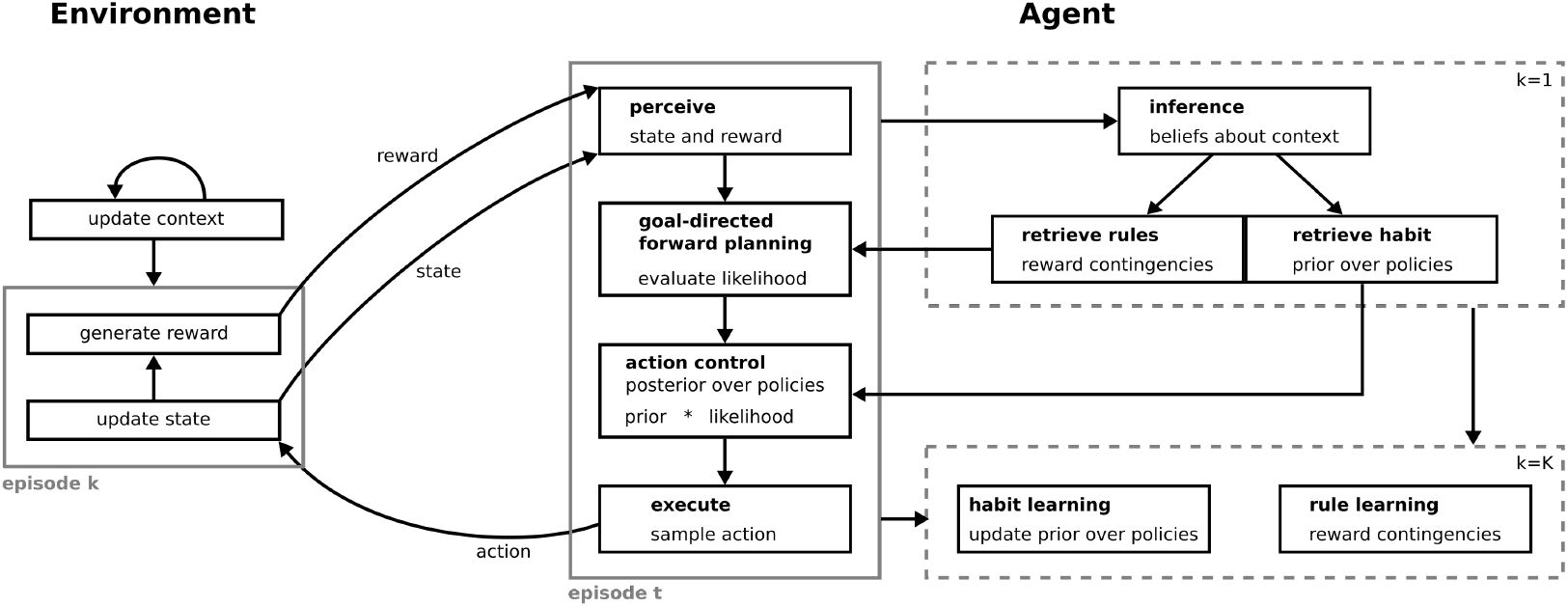
The agent in interaction with its environment. The environment (left) is modeled as a hierarchical partially observable Markov decision process (see Section 2.1). On the lower level, the time evolution of the environment is structured into episodes of length *K*. Here, the states of the environment evolve dependent on the previous state and action chosen by the agent. Given the state and the reward generation rules, some reward or no reward is distributed in each time step *k* of an episode. On the higher level, there is a slowly evolving context which determines the current rules of the environment, namely the reward generation rules, i.e. outcome contingencies. The agent (right) uses its generative model (see Section 2.2 and Figure 2) to represent the dynamics of the environment, and to plan ahead and select actions. At the beginning of each episode (*k* = 1), the agent infers the current context (box in the top right) based on previous rewards and states, and retrieves the learned reward generation rules and the prior over policies for this context. In each time step *k* of an episode (middle box), the agent perceives a new state-reward pair and uses the Markov decision process plan ahead in a goal-directed manner. It evaluates the posterior over policies by combining the control contributions of the prior with the goal-directed control contributions of the likelihood. To execute an action, the agent samples from this posterior. This process repeats until the last time step *k* = *K*, where the agent updates the prior based the chosen policy, and the reward generation rules based on the observed state-reward pairs (bottom right box). This updating is done in a context-specific manner so that prior and contingencies are updated proportionally to the inferred probability of having been in a context during the past episode.

In the second time step *k* = 2 of the *t*-th episode, the environment updates its state to a new state **s**_2_, in accordance with the context transition rule 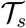, depending on the previous state **s**_1_ and the chosen action **a**_1_. Given the new state and the current context, a new reward **r**_2_ is distributed. The agent once again perceives the state and reward and chooses a new action **a**_2_.

This process is iterated until the last time step *k* = *K* of the episode is reached. In between the last time step of the current episode *t*, and the first time step of the next episode *t* + 1, the context is updated to a new context **c**_*t*+1_ in accordance with the transition rule 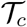. Importantly, the context is an abstract, hidden (latent) state, which determines the current outcome rules of the environment. The context cannot be directly observed by the agent but only inferred from interactions with the environment. We chose this setup because in animal experiments the switch to the context of an extinction phase is typically not cued. Our assumption here is that an agent represents different environments with different rules as different contexts. As in daily life, rule changes might not be directly cued which makes it necessary to model uncertainty about context. This is in line with recent experiments and modelling work which demonstrated that humans and animals implicitly learn different outcome contingencies as different contexts, even when they are not cued (Palminteri et al., 2015; Gershman et al., 2010; Wilson et al., 2014).

Note that this implementation effectively constitutes a hierarchical model on two different time scales: The episodes on the lower level, where states evolve quickly, i.e. in every time step, and the contexts on the higher level, which evolve more slowly, only every *K* time steps.

### 2.2. The generative model

To a participant or an artificial agent, this generative process is not directly accessible. Instead, the agent has to maintain a representation of this process, which is called the generative model. For the purpose of our model, we will assume that the agent knows which quantities are involved: It knows that there are states and that the possible states it could be in are summarized in the set 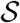. It also knows all possible rewards in 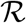, and all possible contexts in 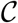.

Furthermore, we assume that the principled structure of the environment is known to the agent: It knows that (i) state transitions depend on the previous state and the action chosen, (ii) reward generation depends on the current state and context, and (iii) the environment is partitioned into episodes, where the context is stable within but may switch between episodes. These causal relationships in the generative model are shown in Figure 2. Within an episode, we assume without loss of generality that the agent does not represent single actions, but sequences of actions (policies)

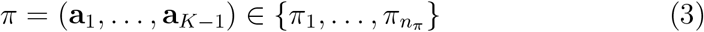

where a policy consists of *len*(*π*) = *K* − 1 actions because actions are executed in between time steps and an action at time step *K* would therefore have no effect.

**Figure 2:**
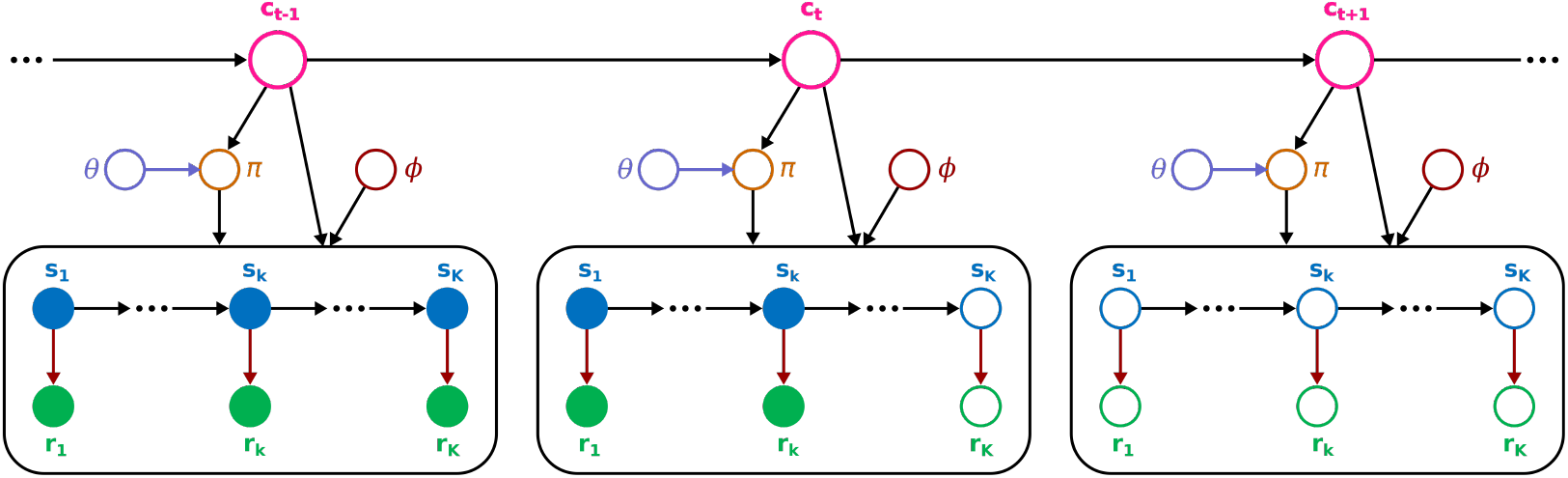
A graphical model depicting conditional dependencies between variables in the generative model. Empty circles indicate latent, unobservable variables and filled circles indicate known, observed variables, and arrows indicate statistical dependencies, where colored arrows indicate that these dependencies are learned by the agent. The model here is a hierarchical model, with the contexts **c**_*t*_ on the higher level of the hierarchy, and the episodes (black boxes) on the lower level of the hierarchy. In the current episode *k* (middle box), the agent starts at in some state **s**_1_ (blue), and receives a reward **r**_1_ (green) according to the current outcome rules (red downward arrows). The agent’s knowledge about the current rules is represented by the parameters *ϕ* (red). The agent then chose some action **a**_1_ in accordance with a policy *π* (brown). For the next time step *k* = 2, the agent transitions to a new state **s**_2_ (arrow to the right), dependent on the policy *π* it followed (downward arrow from *π*), and a new reward **r**_2_ is distributed. This process repeated until the agent reached the current time step *k*. Viewed from here, all future states and rewards are unknown and, so far, unobserved variables, which the agent will infer during its planning process and evaluate if they lead to desirable outcomes. Based on this evaluation of the policies *π* and the prior over policies parameterized by *θ* (lilac), the agent can now choose a new action **a***_k_*. On the higher level of the hierarchy, there are the latent contexts **c**_*t*_ (pink), which evolve more slowly (arrows to the right). They also determine which outcome rules are currently in use (downward right tilted arrow), and which prior over policies is being learned (downward left tilted arrow). The prior over policies is parameterized with the parameters *θ* (lilac), whose influence on the policy is also subjected to learning (lilac arrow to the right). We furthermore show the previous context **c**_*t*−1_ and the next context **c**_*t*+1_, which encode the previous episode (left box) and the next episode (right box), respectively.

Additionally we assume that the agent has the correct representation of the state transition rules 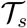. In other words, the agent knows which consequences its own actions will have. In contrast, we assume that an agent does not know the reward probabilities associated with each state and how they depend on the context. Instead, the agent represents those probabilities as random variables

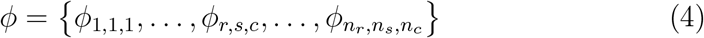

which will have to be inferred.

Importantly, we propose that the agent learns context-dependent habits as a context-dependent prior over policies. It represents the parameters of this prior as latent random variables as well

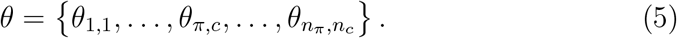

Formally, we write the causal structure of the agent’s generative model as

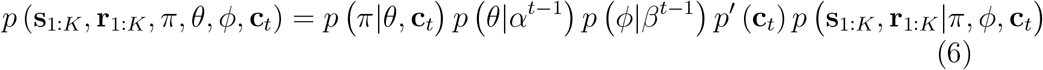

where

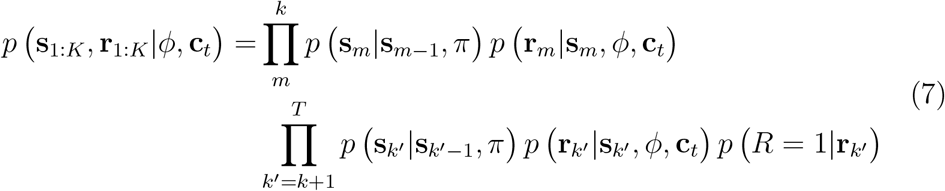

is the agent’s representation of the *k*-th episode, in which it is at time step *k*. This is an effective partition of states and rewards into past observed states **s**_1:*k*_ and rewards **r**_1:*k*_ and unknown future states **s**_*k*+1:*K*_ and rewards **r**_*k*+1:*K*_. The past states and rewards have been observed and are therefore known exactly to the agent. Conversely, the future states and rewards are unknown and are therefore latent variables which will have to be inferred. Note that this is an exact representation of the graphical model in Figure 2.

We use the following distributions to define the generative model:

- The policies *π* are represented by a categorical distribution

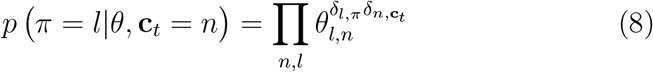

where *δ_i,j_* is the Kronecker delta.
- The latent parameters of the prior over policies *θ* are distributed according to the respective conjugate prior, a product of Dirichlet distributions

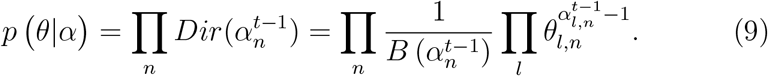
- The so-called concentration parameters 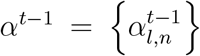 are pseudo counts of the Dirichlet distributions. They encode how often an agent has chosen a policy in a specific context up until the previous episode *t* − 1, and therewith shape the prior over policies.
- The rewards **r**_*k*_ are distributed according to a conditional categorical distribution

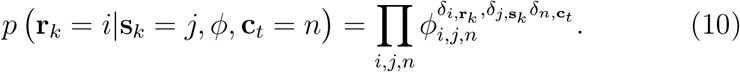
- As above, the latent parameters *ϕ* are distributed according to the product of conjugate Dirichlet priors

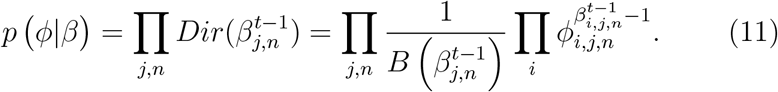
- The concentration parameters 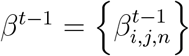 are pseudo counts of the Dirichlet distribution. They encode how often the agent saw a specific reward in a specific state and context up until the previous episode *t* − 1. Therewith they represent the agent’s knowledge about the reward generation rules, i.e. contingencies.
- The states are distributed according to a conditional categorical distribution

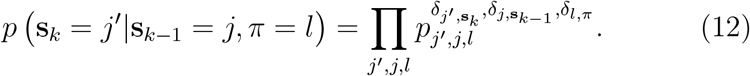 We will fix the parameters *p_j′,j,l_* to the true (deterministic) state transitions 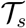 in the generative process.
- The contexts are distributed according to a categorical distribution *p′*(**c**_*t*_). We define this as a predictive prior *p′*(**c**_*t*_) = *p* (**c**_*t*_|**s**_1:*k*−1_, **r**_1:*k*−1_) based on observed past states and rewards. Note that it also includes the agent’s expectation of temporal stability of its environment. Specifically, we assume all contexts have the same temporal stability and change equally often.
- The agent’s preference of rewards is represented by *p*(*R* = 1 |**r**_*k′*_), using a dummy variable *R*, see (Solway and Botvinick, 2012). High values of the probability distribution mean high preference for a particular reward, while low values mean low preference.

After having set up the generative model, we will now show how the agent, based on this model, forms beliefs about its environment and selects actions. To describe action evaluation and selection, we will follow the concept of planning as inference (Attias, 2003; Botvinick and Toussaint, 2012) and active inference (Friston et al., 2015, 2016; Schwöbel et al., 2018). Critically, this means that, apart from forming beliefs about hidden variables of the environment, actions or policies are also treated as latent variables that can be inferred.

### 2.3. Approximate posterior

When an agent infers hidden variables of its environment, such as the context, or future states and rewards, it needs to calculate the posterior

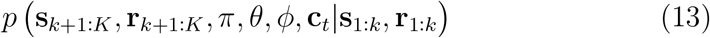

over these hidden variables using Bayesian inversion. Intuitively, this means asking the questions: What context am I most likely in, given I was in these states and received those rewards? What states will I visit in the future, and what rewards will I receive, given I have been in these states in the past and received those rewards? What are the most likely outcome rules that have generated rewards from states? To ensure analytical tractability and low computational costs, we will use variational inference as an approximate Bayesian treatment of the inference process.

Variational inference makes the inference process analytically tractable by replacing the computation of the true posterior with a simpler approximate posterior. In our case we will express the approximate posterior as

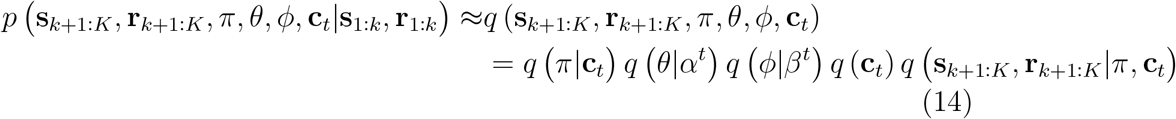

where we use belief propagation based on the Bethe approximation within a behavioral episode

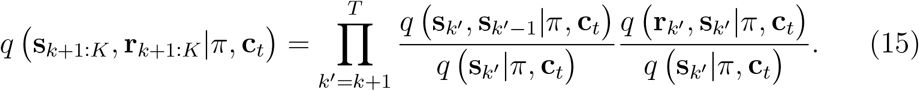

This is well motivated because within an episode, states and rewards critically depend on each other so it is sensible to use an approximation which captures these dependencies.

Outside of an episode, statistical dependencies may be averaged out, so that a mean-field approximation is sufficient to approximate the posterior. Specifically, we will use forward mean-field belief propagation, to obtain an agent’s beliefs based on the observed states and rewards. The posteriors of all random variables will be distributed the same way as in the generative model: states, rewards, policies, and context follow a categorical distribution; while their parameters *θ* and *ϕ* follow a Dirichlet distribution. These come out naturally from calculating the update equations (see Appendix).

### 2.4. Update equations

The marginal and pairwise approximate posteriors can be analytically calculated at the minimum of the variational free energy, see e.g. (Bishop, 2006; Yedidia et al., 2003). These posteriors are typically called beliefs, as they encode the agent’s beliefs about the hidden variables in its environment. We will now show the update equations resulting from the free energy minimization. These equations implement the agent’s information processing: how it forms beliefs about the hidden variables in its environment, how it learns, plans, and evaluates actions. An illustration of this process is shown on the right side of Figure 1.

At the beginning of time step *k* in the *t*-th episode, the agent perceives the state **s**_*k*_ of its environment, and receives a reward **r**_*k*_. It uses this co-occurrence of state and reward to infer the current context and to update its beliefs about the reward generation rules. The posterior over context is estimated as

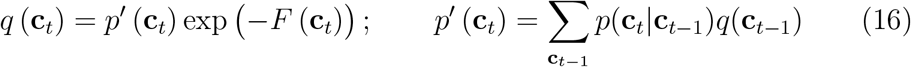

where *p′*(**c**_*t*_) is a predictive probability for contexts given the beliefs previous episode and the transition probabilities *p*(**c**_*t*_|**c**_*t*−1_), and *F*(**c**_*t*_) is the context-specific free energy. The free energy term *F*(**c**_*t*_) encodes the approximate surprise of experienced rewards, states, and the agent’s actions in different possible contexts (see Appendix). The more expected the rewards and actions are for a context, the lower this free energy, and the higher the posterior probability which the agent assigns to this context. As a result, an agent will infer to be in a stable context as long as rewards and actions are as expected, while it will infer a context change if outcomes and actions are unexpected. Note that, initially, before encountering any context, the prior over contexts *p′*(**c**_1_) cannot be set to be uniform. It needs to have a bias towards one of the contexts, so that the agent knows to associate the experienced reward contingencies with the respective context. Which context is assumed to come first is not important, but we found that the agent’s (intuitive) belief that it is most likely in some context is essential for the learning process.

The posterior beliefs about the reward probabilities are again a product of Dirichlet distributions, whose parameters are updated as

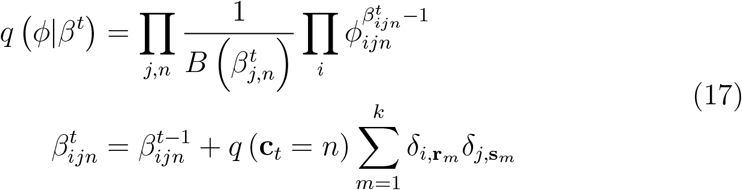

which corresponds to updating pseudo counts 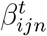. The pseudo counts help keep track of how often the agent has seen a specific reward *i* in a specific state *j* and context *n*. Each time a new reward is generated in a state, these counts are increased by *q*(**c**_*t*_). This way, the counts are high for context with high posterior probability and corresponding observed sequence of reward-state pairs, and low otherwise. At the beginning of a new episode, this posterior will become the new prior, which corresponds to a learning rule in between episodes.

The agent can now use its new knowledge about the rules of its environment to plan into the future and evaluate actions based on their expected outcomes. In order to plan ahead, it calculates its beliefs about future states *q*(**s**_*k′*_) and resulting future rewards *q*(**r**_*k′*_) in the current episode. These beliefs are calculated using belief propagation update rules (see Appendix). If a policy *π* predictably leads to states which yield desirable rewards, as encoded by the outcome preference *p*(*R* = 1|**r**_*k′*_), this policy has a low policy-specific free energy (low surprise) *F*(*π*|**c**_*t*_). The posterior beliefs over policies are computed as

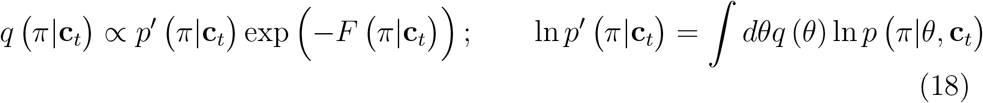

where the free energy corresponds to the log-likelihood in a simple Bayes equation. Importantly, the log-likelihood represents the agent’s goal-directed, value-based evaluation of actions, as it assigns them a value based on predicted future rewards. Additionally, the posterior beliefs contain the prior *p′*(*π*|**c**_*t*_), which assigns an a priori weight to different policies or actions (Doshi-Velez et al., 2010; Todorov, 2009; Friston et al., 2016), where a pronounced prior for a specific policy corresponds to strong habitual control. The agent then samples its next action from the posterior above, which is proportional to the product of the prior times the likelihood. Critically, this leads to an automatic weighting, i.e. arbitration, between goal-directed control (the likelihood) and habitual control (the prior) of the agent’s next action.

At the end of an episode, after having sampled a policy and executed the respective actions, the agent updates its posterior beliefs about the prior over policies

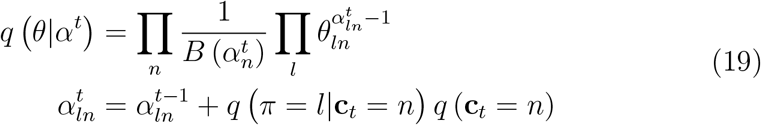

which constitutes habit learning in our model. Here, the pseudo counts 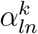 are increased when a policy is chosen in a specific context. After the episode, this posterior becomes the new prior, in order to enable learning across episodes. Note that this implements a tendency to repeat previous actions on one hand, but also to repeat behavior which has been successful in the past. While the prior is independent from the goal-directed evaluation in the likelihood, it is based on which policies were previously chosen. This in turn is influenced by the goal-directed evaluation at the time when they were chosen. In other words, the habit and the outcome rules are learned conjointly. This is an important point because it means that goal-directed control and habit learning are intertwined in a specific way, see also Discussion.

The way the policy pseudo counts 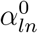 are initialized before the first interaction with any context plays a critical role in how an agent learns a habit. Low initial counts 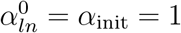 (for every *l, n*) mean that each time a new policy is chosen in a context, the pseudo count increases by a value between 0 and 1 (the posterior over contexts), which increased the count substantially. As a result, the prior over policies becomes fairly pronounced very quickly. In contrast, a high initial count *α*_init_ = 100 means that habits are learned a lot slower, as adding one to this value will have little influence on the prior probability of the corresponding policy. Therefore, we will define a habitual tendency as

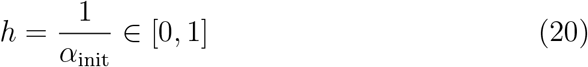

which we will consider a free model parameter with respect to which we will investigate behavioral differences. A high habitual tendency close to 1 will lead to an agent being a strong habit learner and exhibiting fast habit acquisition, while a low habitual tendency close to 0 will lead to a weak habit learning with a low habit learning rate.

### 2.5. Simulation analyses

In this section, we will define quantities which we will use to illustrate our results. Specifically, we will want to investigate how agents infer contexts, using the posterior over contexts *q*(**c**_*t*_), and how agents choose actions, using the marginalized posterior over policies

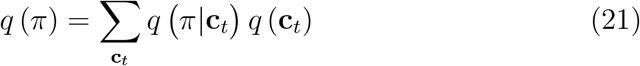

Specifically, to replicate standard results from experimental research, we will report simulations in an environment with two contexts 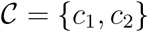 and two actions 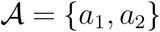. We set episodes to length *K* = 2, so that actions and policies map one to one, which corresponds to a planning depth of 1. We use such short episodes here so that an episode is equivalent to one trial in a habit learning experiment. Nonetheless, it is possible to have longer episodes with increased planning depth in this model, which would endow an agent with the opportunity to learn habits as sequences of actions (see Discussion).

As we have binary random variables, for both contexts and actions we can completely capture the posterior beliefs with a single quantity, the posterior probability of being in second context (*Q_c_* := *q*(**c**_*t*_ = *c*_2_) ∈ [0, 1]) and the posterior probability of selecting the second option (*Q_a_* := *q*(*π* = *a*_2_) ∈ [0, 1]). The posterior probability of being in first context, or selecting first option are obtained as 1 − *Q_c_*, and 1 − *Q_a_*, respectively.

In a similar vein, we also define the likelihood *L_a_* (*t*) := ∑_**c**_*t*__ *q* (**c**_*t*_) exp (−*F* (*π* = *a*_2_|**c**_*t*_))/*Z_c_* of the second option in order to illustrate the agents goal-directed system, and the prior *P_a_* (*t*) := ∑_**c**_*t*__ *q* (**c**_*t*_) *p′*(*π* = *a*_2_|**c**_*t*_) to illustrate how an agent learns habits. The environment will be set to context 1, in a training phase, and switched to context 2 in an extinction phase. When the context switches, the posterior probabilities *Q_c_*, and *Q_a_* should transit from being close to zero, to being close to one, expressing changes in the posterior beliefs as a consequence of the changes in the underlying latent variables. Hence, we assume that the belief trajectory can be fitted with a sigmoid function

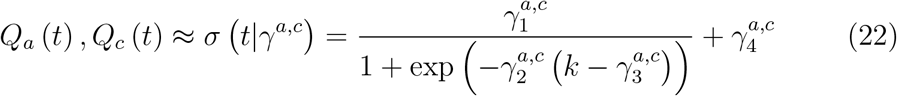

The motivation for this approximation of the trajectory is to determine the trial or episode (*t*\*) at which posterior beliefs *Q_c_*, and *Q_a_* transit from close to 0 to close to 1. The inflection point is specified by the parameters 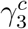 and 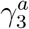, for *Q_c_* and *Q_a_* respectively. We have used the implementation from Python3 SciPy 1.1.0 (Virtanen et al., 2019) of nonlinear curve fitting for this procedure.

We also define a habit strength *H* to quantify the strength of habitual control under different conditions. We define the habit strength as the delay between the actual switch in context of the environment, and the time point at which an agent adapts their behavior. The change in context in our experiment relates to the switch between the training and extinction phases. The time point of adaptation can be interpreted as the trial in which the posterior over actions flips from close to 0 to close to 1. This equates to the inclination point of the sigmoid fitted to the posterior over actions. We define the habit strength as

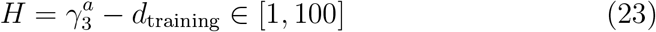

as the difference between the fitted inclination point 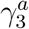 and the training duration *d*_training_. The extinction phase in which we will test for habitual behavior will have 100 trials. As a result, the habit strength can be between 1 and 100, where *H* = 1 indicates that an agent immediately switched its behavior in the first trial of the extinction phase and showed no habitual control, while *H* = 100 means that an agent failed to adapt within the extinction phase and therewith showed full habitual control.

We used the implementation of t-test and ANOVA provided by the Scipy 1.1.0 library (Virtanen et al., 2019). Similarly, we performed the linear regression the implementation of the ordinary least squares (the OLS class) provided in the StatsModels 0.10.1 library (Seabold and Perktold, 2010).

## 3. Results

Having derived the update equations of the prior-based control model, we will now use a series of simulated experiments to show how an artificial prior-based control agent balances its behavior between habitual and goal-directed control. In these simulations, we will use environments where agents are required to adapt their behavior to context switches. We uploaded the code for these simulations to github (https://github.com/SSchwoebel/BalancingControl). In Section 3.1, we will first introduce a task which captures key features of habit learning similar to animal experiments, specifically contingency degradation and outcome devaluation, where we test for habitual behavior in extinction. We will present seven different results:

- We let two exemplary agents perform the task under contingency degradation, show internal properties of the model, and how agents learn habitual behavior (Section 3.2).
- We demonstrate how internal model parameters, like the habitual tendency 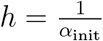, influence the agent’s information processing, behavior, and that an increased habitual tendency increases habit strength after contingency degradation (Section 3.3).
- We show that the acquired habit strength depends on training duration (Section 3.4).
- We show a specific advantage of contextual habit learning, namely that contextual habits allow optimized behavior to be retrieved quickly, when an agent is revisiting a previously experienced context (Section 3.5).
- We show how environmental stochasticity, e.g. highly probabilistic rewards, leads to an over-reliance on habitual behavior and increase habit strength (Section 3.6).
- We introduce outcome devaluation to the task and show that agents exhibit habitual behavior insensitive to contingency degradation and outcome devaluation (Section 3.7).
- We introduce the two-step task (Daw et al., 2011) which has been used to induce habits in humans. We show that the prior-based control agent qualitatively reproduces the behavioral patterns typically found in this task (Section 3.8).

### 3.1. Habit learning task

A common way to experimentally test for habit formation in animal experiments is contingency degradation (Yin and Knowlton, 2006; Wood and Rünger, 2016). Here, an animal is probabilistically rewarded after performing a specific action, e.g. pressing a lever. After a training period, in which the animal learns action-outcome associations and potentially acquires a habit, habitual behavior is measured in an extinction period. The outcome contingencies of the environment are changed, and the lever press does not yield a reward any longer. Conversely, the animal is often rewarded for abstaining from pressing the lever. After this change of contingencies, the strength of habitual control is assessed as the continuation of lever pressing, where a higher habit strength corresponds to more presses. For moderate training durations (~ 50 – 100 trials), the animal will have formed a weak or no habit, and seizes to press the lever rather quickly. For extensive training (~ 500 trials), experiments show that the animal will have formed a strong habit and will continue to press the lever for an extended period of time (~ 50 trials), e.g. (Colwill and Rescorla, 1988; Adams, 1982).

Additionally, for behavior to be classified experimentally as habitual, it must be insensitive to outcome devaluation (Yin and Knowlton, 2006). Here, animals undergo a similar training as in contingency degradation experiments. Then, outcomes are devalued by either satiating the animals, or by associating the reinforcer with an aversive outcome. Afterwards, behavior is again tested in extinction, where a continuing of the lever press is interpreted as evidence for habitual behavior, see e.g. (Adams, 1982). Typically, the strength of habitual behavior also greatly depends on the reinforcement schedule (Yin and Knowlton, 2006), which may be a ratio schedule, where each action leads to a reward with a specific probability, or an interval schedule, where rewards are only distributed after a certain time has elapsed. Interval schedules lead to a greater habit strength and decreased sensitivity to changes in outcome contingencies.

To demonstrate that the prior-based control model can replicate these basic features of habit learning, we approximate the experimental setup of a habit learning experiment in a simplified way, by using a so-called two-armed bandit task, see Figure 3a. This way of modelling the task follows previous modelling studies such as (Daw et al., 2005; Lee et al., 2014) and emulates probabilistically rewarded lever presses of the animal. In the proposed habit learning task, an artificial agent can choose to perform either action *a*_1_, i.e. press a left lever 1, or action *a*_2_ to press a right lever 2. Each lever pays out a reward according to the reward generation rules 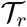, and these probabilities will switch after a certain number of trials, emulating a contingency change, similar to habit learning experiments (Figure 3b). In many habit learning experiments, the animals do not choose between two levers, but rather between pressing a lever or abstaining from pressing, where abstaining is a viable option due to opportunity costs. We approximated opportunity costs of not pressing the lever by introducing a minimally rewarded second choice (lever 2) instead, see also similar approaches taken in previous modelling studies (Daw et al., 2005; Lee et al., 2014; Keramati et al., 2011; Pezzulo et al., 2013; Gershman et al., 2014).

**Figure 3:**
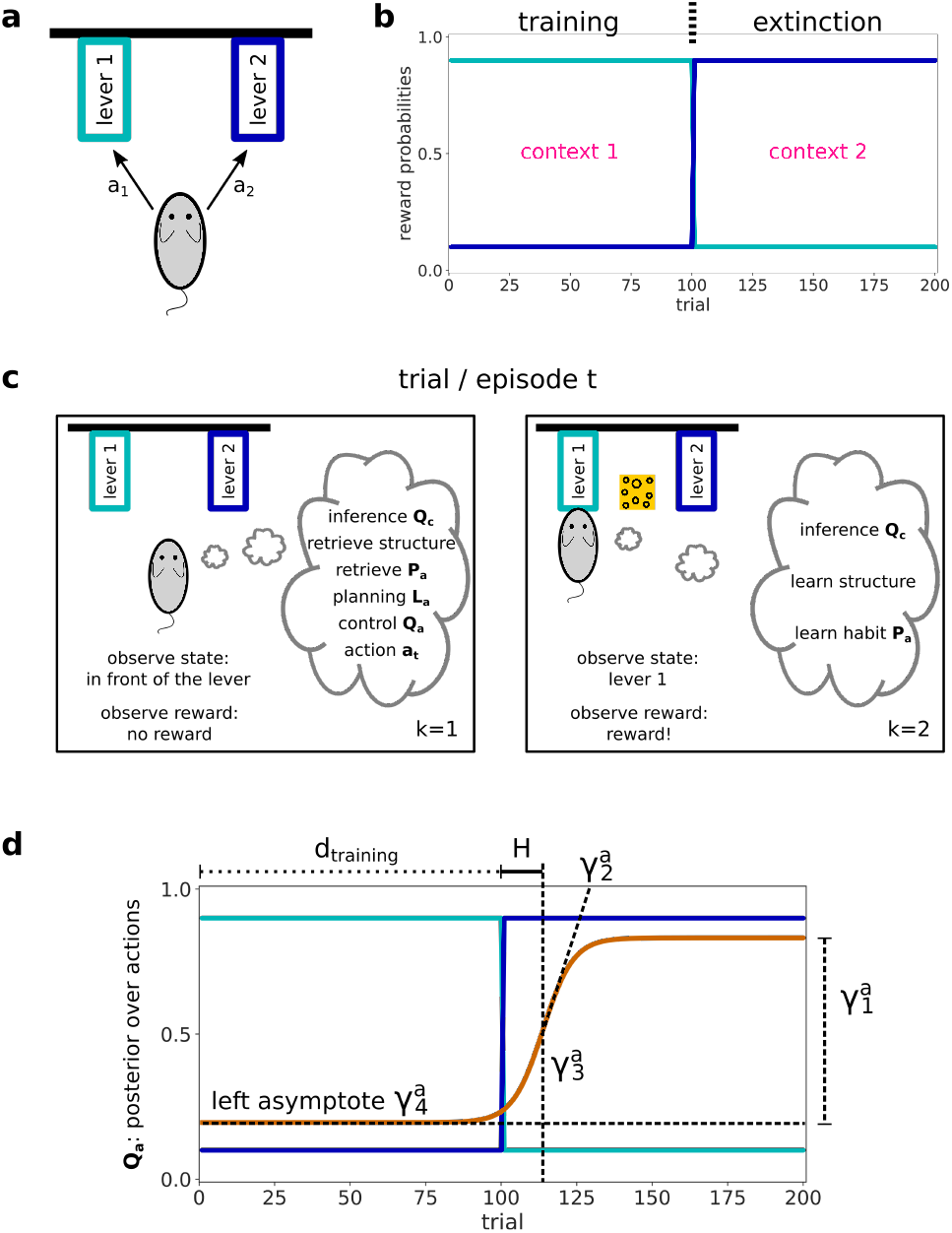
Habit learning task. **(a)** In each trial *t*, the agent can choose between pressing two levers (light and dark blue boxes, lever in black next to the box) and is awarded probabilistically. We model this task as a two-armed bandit task. **(b)** Reward schedule over 200 trials for the two levers. In the training phase, lever 1 yields a reward with *ν* = 0.9 probability, while lever 2 only yields a reward with 1 – *ν* = 0.1 probability. After 100 trials, the reward probabilities switch. The new contingencies are stable for another 100 trials. This second stable period emulates an extinction phase, where we will test the agent’s habit strength by how quickly it is able to adapt its choices. **(c)** An agent solving the task. For the agent, each trial constitutes one behavioral episode. In episode or trial *t*, the agent starts out in the state (position) in front of the two levers in the first time step *k* = 1 of this episode. It observes its state and that there is no reward. The agent can now infer the context *Q_c_* based on its experience in the previous trials. It retrieves the learned outcome contingencies and habit *P_a_* for this context from memory. It uses its knowledge about the reward structure to plan forward and evaluate actions based on the likelihood *L_a_*, where actions which lead more likely to a reward will have a higher likelihood encoding the goal-directed value. The agent combines the likelihood and the prior to evaluate the posterior over actions *Q_a_* and samples a new action **a***_k_* from this posterior, for example action *a*_1_. In between episodes, this action is executed and the agent transitions to the new state, pressing lever 1. At the beginning of the next time step *k* = 2, a reward may be distributed, depending on the action and lever the agent chose. It then updates its context inference *Q_c_* based on the perceived state-reward pair, learns the outcome rules, and updates its prior over actions *P_a_*. This process repeats until the last trial *t* = 200. **(d)** Illustration of the sigmoid function used to analyse the time evolution of the posterior over actions *Q_a_* (see Section 2.5 for details). The *a* as a superscript on the parameters signifies that these are the parameters for the posterior over actions. We define the habit strength *H* as the difference between the inflection point of the posterior beliefs 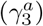 and the trial number at which the context changed *d*_training_.

The habit learning task has two phases (Figure 3b): The first phase is the training phase which lasts *d*_training_ = 100 trials. We will also vary this duration in Section 3.4. Here, lever 1 pays out a reward with *ν* = 0.9 probability, and lever 2 with 1 – *ν* = 0.1. These reward probabilities are kept stable during the training period and the agent learns about outcome contingencies and might form a habit. The second phase is the extinction phase which lasts another 100 trials. Here, outcome probabilities are switched relative to the training phase, and are kept stable for the remainder of the experiment. After the switch of outcome contingencies, we quantify an agent’s habit strength as the number of trials before an agent adapts its behavior and primarily presses lever 2 instead of lever 1, see section ‘Simulation analyses’ in Methods. Note that in our simulations, due to our agent setup, a trial is equivalent to a behavioral episode for an agent, see Figure 3c for an exemplary episode in which the agent interacts with the habit learning task.

This experimental setup emulates the training and extinction phases of a contingency degradation habit learning experiment. It can be transformed into a outcome devaluation experiment by modulating the agent’s preference for outcomes (*p* (*R* = 1|**r***_k_*), see Section 2.2 and Appendix) after the training phase. In order to disentangle these two effects, we will restrict our simulated experiments to contingency degradation in most of the following sections. In the last section, we will show habitual behavior under outcome devaluation.

Note that the two phases of the experiment (Figure 3b) can be viewed as a sequence of two contexts, where in each context one of the two choices returns higher expected reward. Importantly, the agent is initially not explicitly aware how any context is associated with a specific set of outcome rules. Instead, the agent learns to associate the outcome rules it first experiences with the first context. When the contingencies change, it will infer the change and learn to associate the new rules with a second context. The prior-based control model is general enough to be able to deal with an arbitrary number of contexts, and also arbitrary contexts, where e.g. not one lever is clearly the better option. For details, we refer the reader to the Supplementary material.

By design in our experiment, this corresponds to associating contexts with preferable levers. In some habit learning experiments, contexts are cued and habitual behavior is used in response as form of stimulus response association, e.g. (Sage and Knowlton, 2000). In our habit learning task, we do not use a cue to indicate the context to the agent. This is in line with typical animal experiments where the extinction phase is not cued. Instead, the state, i.e. the position of the agent in front of the levers is observable and takes the role of a stimulus.

### 3.2. Habit learning under contingency degradation

In this section, we illustrate, in detail, how agents based on the prior-based control model learn about their environment, form beliefs, acquire habits, select actions, and balance goal-directed and habitual control, see Methods and Figure 1. As the habitual tendency parameter *h* has a strong influence on habit learning and action selection, we will show two exemplary simulations of a an agent with strong (*h* = 1.0) and another agent with a weak (*h* = 0.01) habitual tendency performing the task (Figure 3). In the following, we refer to these two agents as the strong habit learner (*h* = 1.0) and the weak habit learner (*h* = 0.01). Note that, in this section, for didactic purposes, we will describe model behavior on just single instances of two representative agents. This is followed by more thorough simulations, where we also quantify the uncertainty over model variables using multiple experiments for each agent.

When an agent is first put into the task environment, it has no prior knowledge about the outcome contingencies associated with any context, and no prior preference for any actions 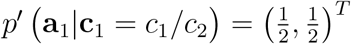, i.e. there is no habit yet. What the agent does know, is that action 1 means pressing lever 1, and action 2 means pressing lever 2, so that it has an accurate representation of the state transition matrices *p* (**s**_*k*+1_|**s***_k_*, **a***_k_*). Furthermore, the agent has a prior over contexts with a bias towards context 1 (see Methods).

In the first trial, the agent has not sampled any reward yet, so it chooses an action **a**_1_ randomly as it does not have any knowledge available to predict the outcome of actions. According to the action chosen, the agent goes to and presses the respective lever, and receives a reward or no reward. At the end of the trial, as this also marks the end of a behavioral episode, the agent updates its prior *P_a_* to increase the a priori probability to repeat this chosen action, and updates its knowledge about the reward structure (see Figure 1 and Figure 3c). As the agent started with a biased prior over contexts, it associates this reward structure with context 1. Hence, the prior bias for context 1 simply reflects agent knowledge that it can be in only one context initially.

At the start of the second trial, the agent infers that it is most likely in context 1 (*Q_c_*), based on its previous experience and its knowledge about the stability of the environment. It retrieves the reward structure and the prior *P_a_* over actions it just learned. The agent can now use this new knowledge about outcome contingencies in the current context to evaluate the likelihood *L_a_*. In order to select an action, it calculates the posterior beliefs over actions *Q_a_* as the product of the prior *P_a_*, which represents habits as an automatic and value-free tendency to repeat actions, and the likelihood *L_a_*, which represents the goal-directed and value-based evaluation of anticipated future rewards (see Eq. 18). The agent then samples an action **a**_2_ from these posterior beliefs about actions, dynamically adjusting the balance between goal-directed and habitual choices. The agent visits and presses the lever it just chose and samples a reward. At the end of this trial and behavioral episode, the agent reevaluates its beliefs about the context *Q_c_*, based on if the new observations still fit to its knowledge about this context. The agent also updates its prior over actions *P_a_*, hence increasing the prior probability of that action being repeated (thereby learning a habit). Similarly, the agent updates its knowledge about the reward structure, based on its beliefs about the context. This update cycle is repeated over all future trials, see Figure 3c and Section 2.4.

Figure 4 shows the resulting dynamics of the relevant agent variables (*Q_c_*, *L_a_*, *P_a_*, *Q_a_*, **a***_t_*) for the strong (left) and weak (right column) habit learner during all 200 trials in the habit learning task. In the training phase, the beliefs over context *Q_c_* converge rather quickly and after about 10 trials, the two agents are certain of being in context 1, see Figure 4a). Figure 4b shows the likelihood over actions *L_a_*, reflecting the expected choice value, that is, the estimated surprise in reaching a goal (observing a rewarding outcome). As the likelihood depends on the learned knowledge about the environment, it takes both weak and strong habit learners around 30 trials to observe enough outcomes before the likelihood converges to a stable value. Figure 4c shows how the prior over actions *P_a_* evolves, i.e. how a habit is learned. Here, the difference between the strong and weak habit learner is obvious: The strong habit learner (left) forms a strong habit quickly (*P_a_* < 0.1) after only 40 trials. This means, the strong habit learner has a very high a priori probability 1 – *P_a_* of choosing action 1 independent of the expected rewards. Conversely, the weak habit learner updates its prior over actions rather slowly (*P_a_* ∈ [0.4, 0.6]). The second to last row (Figure 4d)) shows the posterior over actions *Q_a_*, which is the product of the prior and the likelihood. For the weak habit learner, the prior has little to no influence, as it is close to 0.5, so that the posterior over actions looks similar to the likelihood. For the strong habit learner, the strong prior lets the posterior over actions converge to values close to 1.0 within 40 trials. The agents sample their actions from this posterior probability, which are shown in the bottom row (Figure 4e)). The strong habit learner chooses the action with the higher expected reward more consistently (94% of choices), while the weak habit learner continues to choose action 2 even late into the training period. As a result, the weak habit learner has a significantly lower success rate (80%, *p* = 0.003, two sample t-test on the chosen actions in the training phase of two agents shown here).

**Figure 4:**
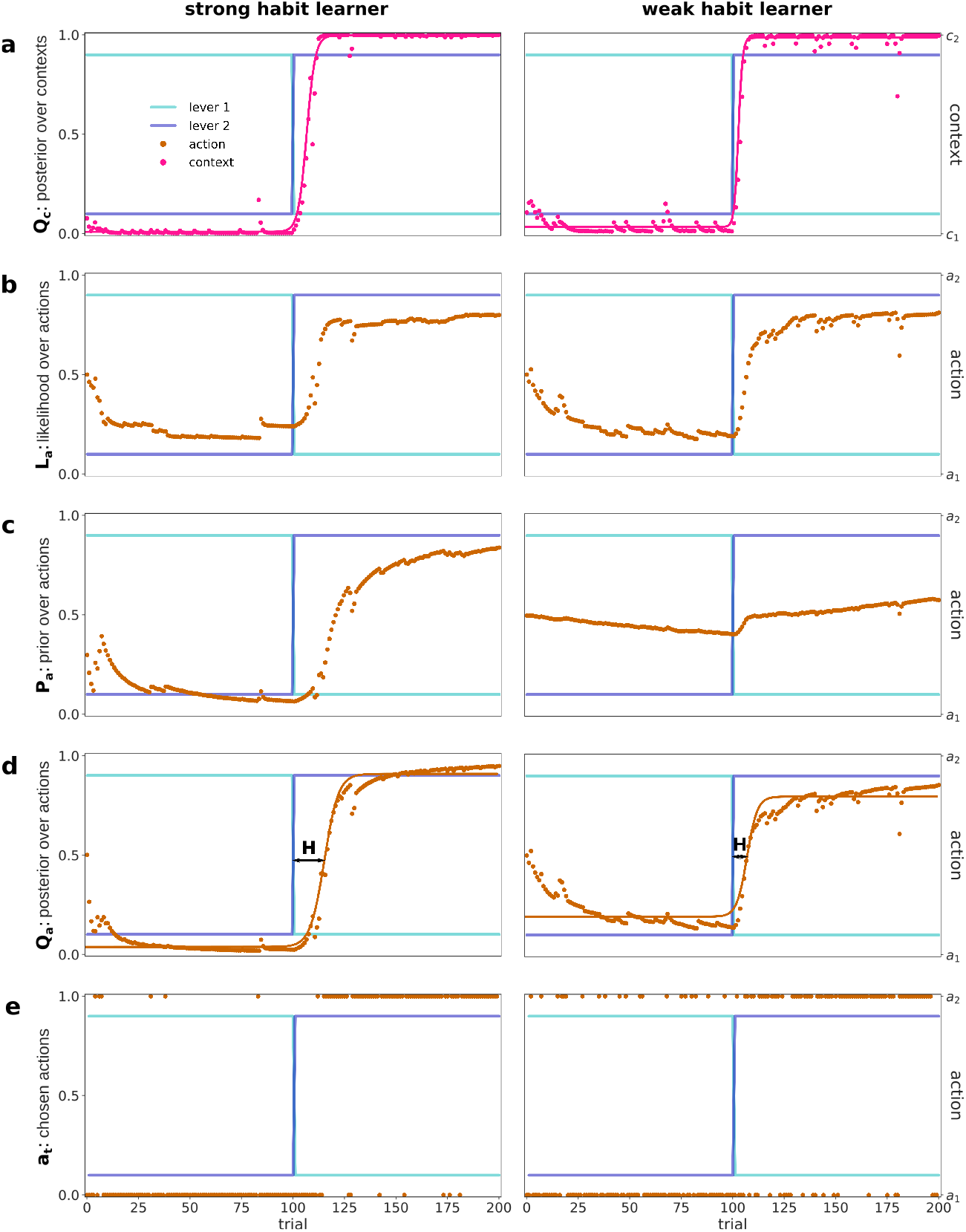
The dynamics of key internal variables of contextual habit learning agents during the habit learning task. The left column shows the dynamics for a strong (*h* = 1.0) habit learner and the right column for a weak (*h* = 0.01) habit learner. **(a)** The first row shows the agent’s inference, the posterior beliefs over contexts *Q_a_*, i.e. the estimated probability of being in context 2. The pink dots are the agents’ posterior beliefs in each trial of the task. The pink solid line is a fitted sigmoid, where its inclination point 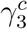 indicates when the posterior changes from representing context 1 to context 2. The light and dark blue lines are the reward probabilities of levers 1 and 2, respectively (see Figure 3). **(b)** The brown dots in the second row show the (normalized) likelihood *L_a_* over actions. The likelihood encodes the goal-directed, expected reward of actions, given the learned outcome contingencies. **(c)** The brown dots in the third row show the prior over actions *P_a_*, which encodes how likely the agent is a priori to select lever 2. **(d)** The fourth row shows the posterior over actions *Q_a_*, which is the product of the prior and the likelihood. The brown dots show the posterior in each trial of the task, and the brown solid line shows a fitted sigmoid, whose inclination point can be interpreted as the trial at which an agent adapts its actions (see Figure 3d). The black arrow shows the resulting habit strength *H* which is the inclination point minus the training trials. **(e)** The brown dots in the bottom row show the chosen actions, which were sampled from the posterior over actions.

In the extinction phase, after the switch in trial 100, the reward contin-gencies become reversed. When continuing to press lever 1, the agents are only rewarded with a probability of 1 – *ν* = 0.1. The lack of expected reward payout produces a prediction error which increases the context-specific free energy (see Section 2.4). This drives the agents to quickly infer that the previously inferred context 1 is no longer an appropriate representation of the environment (see Figure 4a). Instead, the agents switch to believing to be in a new (second) context, and learn reward contingencies and habits for this context. Note that the model is general enough to learn any number of contexts with arbitrary outcome contingencies. For an illustration of this property, we refer the reader to the Supplementary material. The weak habit learner infers the context switch slightly earlier than the strong habit learner, at trials 103 and 107, respectively. In the prior-based control model, the agents’ context inference not only depends on surprising outcomes but also on the agents’ own actions (see Section 2.4). The strong habit learner behaves highly consistently, even after the switch, and therefore is delayed in its context inference, relative to the weak habit learner. Note that the time point of this switch in beliefs was measured as the inflection point of a sigmoid fitted to the beliefs over time (a; solid line), see 2.5 and Figure 3d for a detailed explanation of how we used the parameters of the sigmoid.

Following context inference, the agents learn the new reward contingencies (see Figure 4b) and new habits (see Figure 4c) for context 2. Since this learning takes place after the context inference step, the posterior over policies is updated with a delay with respect to the context inference. As the agents sample their actions from the posterior, we can measure the trial at which they adapt their actions to press mostly lever 2 as the inflection point of the posterior. As with the posterior over contexts, we fitted a sigmoid (solid lines in Figure 4d) to calculate the time point of action adaptation, see Section 2.5 and Figure 3d.

In the following, we will call the time point (in trials) of action adaptation after the contingency change the habit strength, see 2.5. A value of 1 corresponds to the lowest possible habit strength, while a value of 100 means that an agent completely failed to adapt its behavior. This quantification is in line with the animal literature, where the amount of habitual behavioral control is measured by how often animals continue to choose the previously reinforced action after contingency degradation. As expected, the strong habit learner adapts its behavior later than the weak habit learner, at trials 116 and 107, respectively. This means the strong habit learner has a habit strength of 16 and the weak habit learner of 7.

The actions after the contingency switch in Figure 4e reflect this quantification of habit strength. The strong habit learner continues to choose lever 1 for around 10 trials, before it adapts and mostly consistently chooses lever 2 after 20 trials. The weak habit learner adapts earlier, but behaves less consistently and requires a longer transition period where both actions are chosen. However, due to the faster adaptation, in the first 15 trials after the switch, the weak habit learner exhibits a higher performance (chooses lever 2 in 47% of trials) than the strong habit learner (7% of trials, *p* = 0.012, two sample t-test on the actions in the first 15 trials after the switch).

The strong habit learner is able to recover its performance in the remainder of the extinction phase, where the task context is once again stable. Here, it not only learns the new reward contingencies, but a strong prior for action 2 (Figure 4c), so that it is again able to choose lever 2 more consistently, relative to the weak habit learner (92% vs 78%, *p* = 0.01, two sample t-test on the actions in trials 116 – 200).

In summary, we found that a more pronounced prior causes a stronger habit, as measured by the number of trial in the extinction phase before behavior is adapted. Critically, the mechanism is that a strong prior (Figure 4c) increases the certainty in the agent’s posterior over actions (Figure 4d) and thereby its selection of the action (Figure 4e) with the higher expected reward. We found that as long as the environment is stable, the strong habit learner chooses the more rewarding option more reliably. This is the case in the training phase until the switch, and – after a brief adaptation period – after the switch. The strong habit learner exhibits less optimal behavior, in terms of obtained reward and relative to the weak habit learner, only immediately after the switch. This indicates that being a strong habit learner is useful for an agent, as long as contexts do not switch too often.

In addition, note that the effect of an increased certainty in action selection caused by the prior over actions is similar to a dynamic adjustment in decision temperature. Here, we did not use a decision temperature in our decision rule, as would be usually done in modeling noisy behavior (of participants), see Methods. Rather, we let the influence of the prior take this role. In the proposed context-specific model, this seems well motivated as the prior is learned conjointly with the reward contingencies, and indirectly reflects which behaviors have been successful in the past. This means that, in the proposed model, learned habits express themselves not only as an a priori preference for an action, but also as a dynamic adjustment of a decision temperature.

### 3.3. Habitual tendency increases habit strength

To generalize the effect of the habitual tendency on an agent’s beliefs and behavior, we analysed agents with different values of the habitual tendency *h*, where we repeated simulations for each value 200 times, see Figure 5. The results confirm the conclusions drawn in the previous section: (i) All agents, independent of habitual tendency infer the context change quickly (within the first 5 trials after the switch), where strong habit learners infer the switch slightly later (*p* = 0.01, linear regression on the median values). (ii) Behavioral adaptation is at least 5 trials delayed compared to context switch inference. We find that acquired habit strength increases with the habitual tendency of an agent (*p* < 0.001, linear regression on the median values).

**Figure 5:**
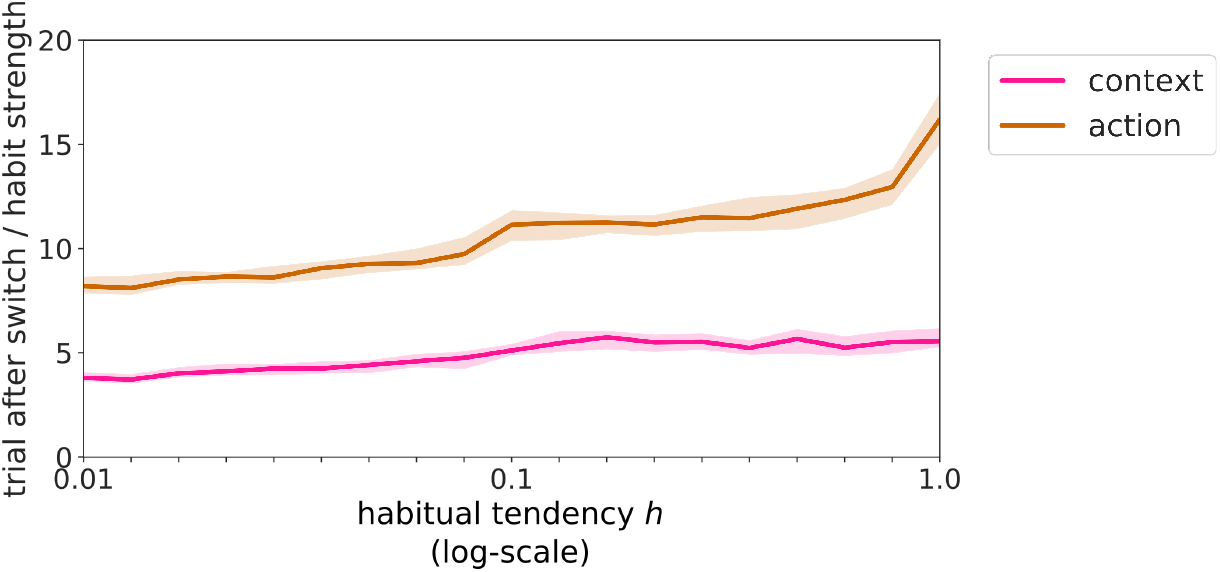
Habit strength as a function of the habitual tendency. For values of habitual tendency between 0.01 and 1.0, we plot the time points (in trials) of an inferred switch in context (pink solid line) and the habit strength (brown solid line). We measure habit strength as the time point of action adaptation after the switch, see Methods. For each habitual tendency value, we plot the median of 200 simulated runs, where the shaded areas represent the confidence interval of 95% around the median. We found a significant correlation between habitual tendency and habit strength (*p* < 0.001) and between habitual tendency and context inference (*p* = 0.01). The x-axis is logarithmically scaled.

### 3.4. Training duration increases habit strength

Here, we show that the prior-based control model is able to capture experimental findings that acquired habit strength depends on the amount of training a participant received. To test this, we simulated agents in the same habit learning task as above (see Figure 3) but now vary the length of the training phase *d*_training_ before the extinction phase.

In Figure 6 we plot the habit strength (see Methods) for three representative agents with different habitual tendencies (strong (*h* = 1.0), medium (*h* = 0.1), weak (*h* = 0.01)) as a function of training duration. For moderate training period durations (*d*_training_ ≤ 100 trials), agents develop a relatively low habit strength and adapt their behavior rather quickly, within 20 trials. Although the differences are small for moderate training lengths, we find, as in the previous section, a significant correlation between habit strength and habitual tendency (*p* < 0.001, linear regression on the median values).

**Figure 6:**
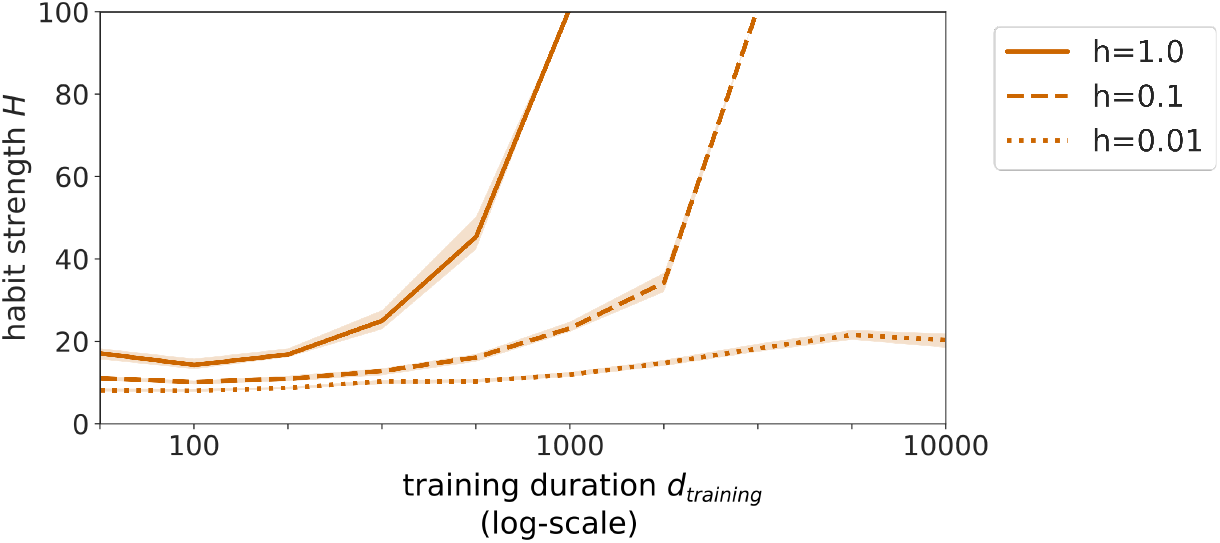
Habit strength as a function of training duration *d*_training_. The x-axis is scaled logarithmically. The solid line represents a strong habit learner with a habitual tendency of *h* = 1.0, the dashed line a medium habit learner with *h* = 0.1, and the dotted line a weak habit learner with *h* = 0.01. The lines show the medians estimated over *n* = 200 repeated simulations for each level of the habitual tendency *h*. The shaded area shows the 95% confidence interval. A habit strength of 100 means that the posterior choice probability *Q_a_* remains smaller than .5 during the entire 100 trials of the extinction phase.

For longer training durations, habit strength is generally increasing. For very long training durations, both the strong and medium habit learner fail to adapt their behavior within the extinction period of 100 trials. The strong habit learner cannot adapt for a training duration *d*_training_ ≥ 1000, and the medium habit learner for a training duration greater 5000. The weak habit learner exhibits only a slight increase in habit strength as a function of training duration. This is to be expected, as the weak habit learner learns habitual control so slowly, that it is close to being purely goal-directed.

In summary, these results stress the role of learning a prior over actions, where we interpret a strong prior as the representation of a habit, see e.g. Figure 4d. The longer the training period, the more pronounced the prior of a specific action will be, while the likelihood stabilizes after contingencies have been learnt properly (around 40 trials). Therefore the prior’s influence on context inference and action adaptation increases with longer training periods, so that agents choose the previously reinforced action longer and longer in the extinction phase. The exact training duration at which adaptation starts to be delayed and fail depends on an agent’s individual habitual tendency, where a higher tendency leads to a fail in adaptation for shorter training periods. This is in line with the literature on moderate and extensive training, where extensive training leads to increased habit strength (Seger and Spiering, 2011).

### 3.5. Retrieval of previously learned context-specific habits

So far, we have assessed how habits can be represented as a prior over policies, where this prior is learned in a context-specific fashion. Here, we show a specific advantage of this context-specificity: The agent can recognize a previously experienced context by the associated contingencies and retrieve its habit (i.e., prior over actions) and learned reward generation rules for this context (Bouton and Bolles, 1979). As the prior implements a tendency to repeat actions, and actions were chosen according to their usefulness (i.e., likelihood of being chosen, see Fig. 1), habits encoded in the prior-based control model represent which behavior is advantageous in a specific context. Therefore, recognizing the context and reusing previously established priors corresponds to a retrieval of previously learned optimal behavior, i.e., habits.

In Figure 7, we show the design of an ABA renewal experiment Bouton and Bolles (1979); Gershman et al. (2010); Nakajima et al. (2000); Bouton et al. (2011), which is an extension habit learning task. As before, we first let agents experience the two contexts for 100 trials each, and call this the learning phase of the experiment. Critically, there is an, additional phase, the renewal phase, where we place agents again into context 1 for 100 trials. In the first trial of this renewal phase, we induce maximal uncertainty about the context by setting the agents’ prior over contexts to *p* (*c*_201_) = (0.5, 0.5)*^T^*. Here, we wanted to emulate a situation where an agent knows there is a context change, but not to which context, akin to a mouse being taken out of its home cage into the experimental setup. Apart from setting the prior over contexts, we did not cue a context change in this experiment. If we had kept the prior over contexts as the old posterior from the last trial of the learning phase, we would induce habit effects where agents delay adaptation for the reasons discussed in the previous sections. To compare ‘experienced’ agents with agents that have not learned yet context 1, we implement ‘naive’ agents, as in Section 3.2.

**Figure 7:**
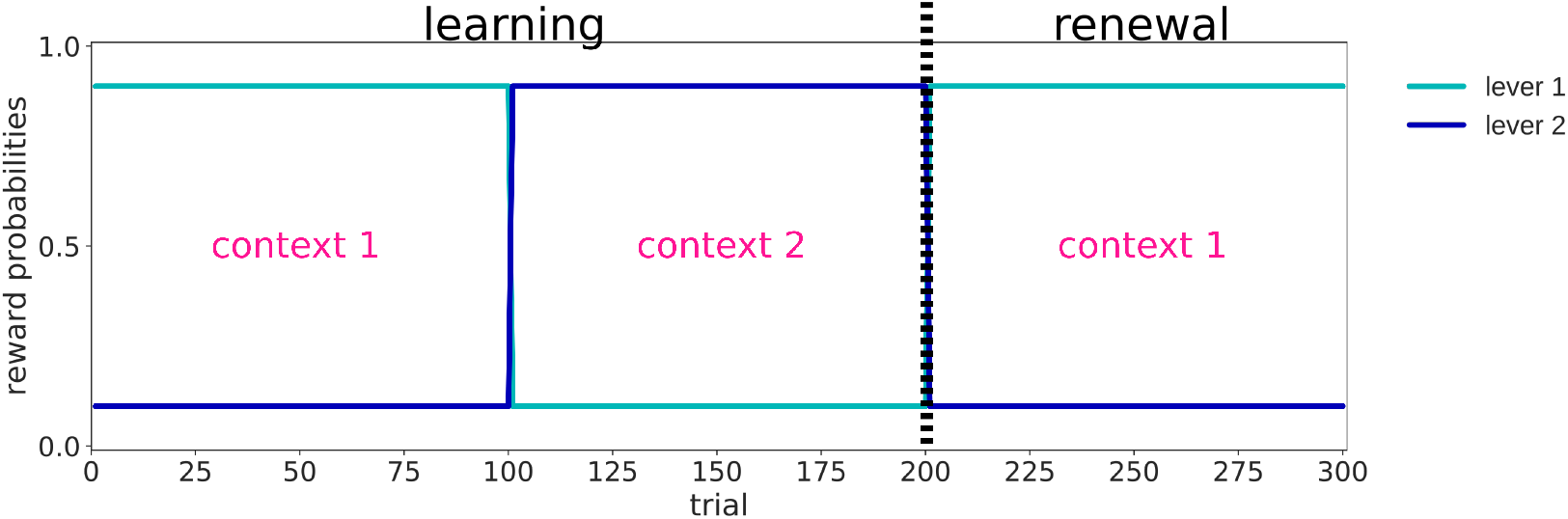
The ABA renewal experiment. A 300 trial experiment consisting of a learning phase (equivalent to the whole habit task, i.e. the conditioning and extinction phase, see Figure 3) with 200 trials, and a new, additional habit renewal phase with 100 trials. The light blue line shows the probability of lever 1 paying out a reward, and the dark blue line shows the probability of lever 2 paying out a reward. The vertical dashed line indicates the switch from the learning to the renewal phase. In the renewal phase, the agent revisits context 1, where outcome contingencies are exactly the same as in the first 100 trials of the experiment.

To quantify the advantage of the retrieval of previously learned context-specific behavior in the renewal phase, we first measured how long it takes a naive agent to converge to a stable beliefs level about context 1 in the learning phase (Figure 8a). To evaluate the convergence times to a stable knowledge for naive agents, we fit again a sigmoid to the posterior over contexts and actions in the learning phase (as in Figure 4, see also Methods). We interpret the left asymptote 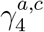 of the sigmoid as the stable level of knowledge the agents eventually reach. We calculate the convergence time as the trial in which the posterior crosses the left asymptote for the first time. We compare this duration to how long experienced agents take to recognize the known context 1 in the renewal phase and reuse their previously learned behavior. To compute convergence times for the experienced agent, we determined the first trial in the renewal phase where the posterior is lower than the left asymptote which was fitted for the learning phase.

**Figure 8:**
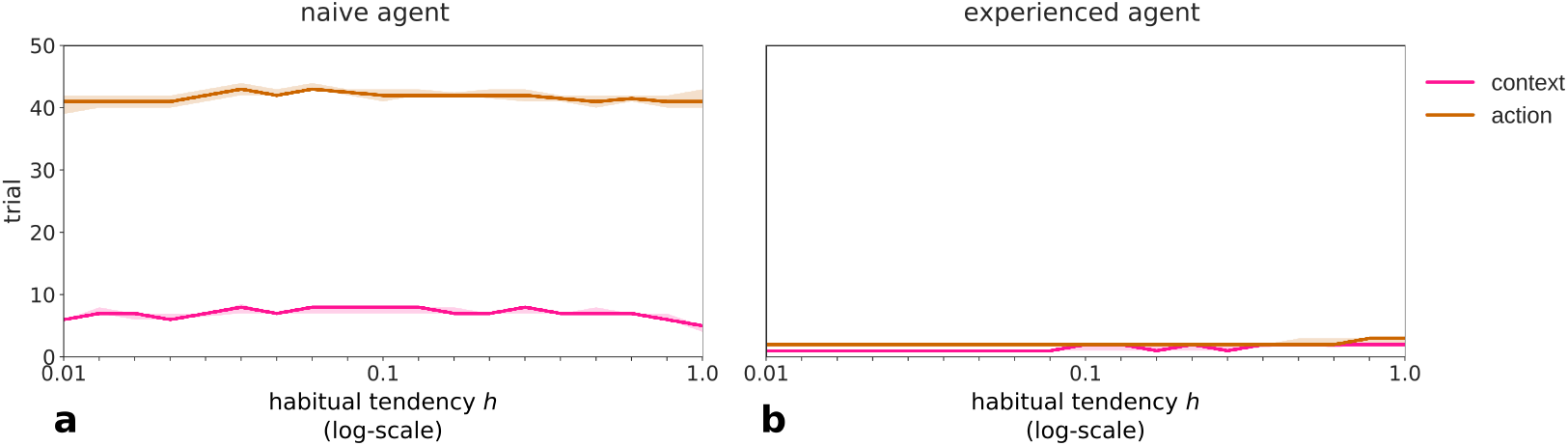
Convergence times of the posterior over contexts (pink) and posterior over actions (brown) in naive (a) and experienced (b) agents as a function of habitual tendency. The shaded areas indicate a confidence interval of 95%. a) Convergence times of the posterior beliefs in naive agents who visit context 1 for the first time, see main text how convergence times were quantified. A naive agent takes around 7 trials to converge to stable beliefs about its context. It takes around 40 trials to converge to a stable posterior over actions, indicating the time it takes to learn a stable representation of the action-outcome contingencies for this context. b) Convergence times of the posterior beliefs in naive agents who visit context 1 for the second time. An experienced agent takes 1 to 2 trials to recognize it is in the known context 1. It almost instantly retrieves its knowledge about outcome contingencies and its habit for this context, and thereby its posterior over actions, so that the action adaptation happens maximal one trial later.

These convergence times, as a function of habitual tendency, are shown in Figure 8 for both the naive and the experienced agents. We discussed the initial development and convergence of the posteriors shown in Figure 4 for single runs of agents in Section 3.2. The results here are a quantification of these for different habitual tendencies using 200 runs each. Naive agents (see Figure 8a) are able to achieve a stable level of knowledge for the context in around 8 trials, if they have a low habitual tendency (e.g. 0.02), and in around 5 trials, if they have a high habitual tendency (1.0). As discussed above, context convergence time are faster for higher habitual tendency, because these depend partially on the agent’s own more consistent behavior. Action convergence times mainly depend on learning the outcome rules and the resulting likelihood, which takes, for the naive agent, with around 40 – 45 trials a lot longer than context inference. We find that these times are not influenced by an agent’s habitual tendency.

For experienced agents, both, recognition of the known context, as well as reusing the old outcome rules and habits, happens almost instantaneously, within the first 3 trials of the retrieval phase, see Figure 8b. As a consequence of these faster convergence times, experienced agents choose the optimal lever more often in the retrieval phase than in the first half (context 1) of the learning phase (94% vs 87%, *p* < 0.001, two-sample t-test, averaged over all habitual tendencies). In addition, we find that agents continue to learn outcome contingencies and habits during the renewed exposure to context 1 (data not shown). Importantly, in terms of behavior, for both the naive and experienced agent, the percentage of choosing the optimal action increases with habitual tendency (naive: *p* = 0.001; experienced: *p* = 0.036; linear regression on the median values). This finding provides another hint that being a strong habit learner might be advantageous if one’s environment is mostly stable except for sudden switches to already known contexts, see also Discussion.

### 3.6. Environmental stochasticity increases habit strength

In this section, we examine how environmental stochasticity, namely the probability of observing a reward, interacts with the habit learning process (DeRusso et al., 2010). We again let artificial agents perform in the habit learning task (see Figure 3). We varied the probability of receiving a reward *ν* in both the training and extinction phases from *ν* = 1.0 (completely deterministic) to *ν* = 0.6 (highly stochastic, where a 0.5 probability would mean that outcomes are purely random). In the extinction phase, lever 1 has probability *ν* to pay out a reward, while lever 2 pays out a reward with a probability of 1 – *ν*. These probabilities are reversed in the extinction phase.

Figure 9 shows the habit strength, measured in the extinction phase as a function of environmental stochasticity 1 – *ν*. As before, we used three agents with different habitual tendencies (strong (*h* = 1.0), medium (*h* = 0.1), weak (*h* = 0.01)). In a fully deterministic environment (1 – *ν* = 0), all three agents have a similarly low habit strength (below 10). The agents infer the context switch immediately (not shown) and adapt their behavior shortly after. When the reward probability is *ν* = 0.9 and the stochasticity is 1 – *ν* = 0.1, we find habit strengths between 7 and 15, which replicates the result shown in Figure 5. For more stochastic rewards, we find that for all three agents the habit strength increases with stochasticity, until they fail to adapt within the extinction phase. In addition, one can see that the habit strength is higher, the higher the habitual tendency of the agent is (*p* < 0.03 for a ANOVA on parameters of fitted exponential functions), and the exact amount of stochasticity agents can handle before they fail to adapt depends on the agent’s habitual tendency.

**Figure 9:**
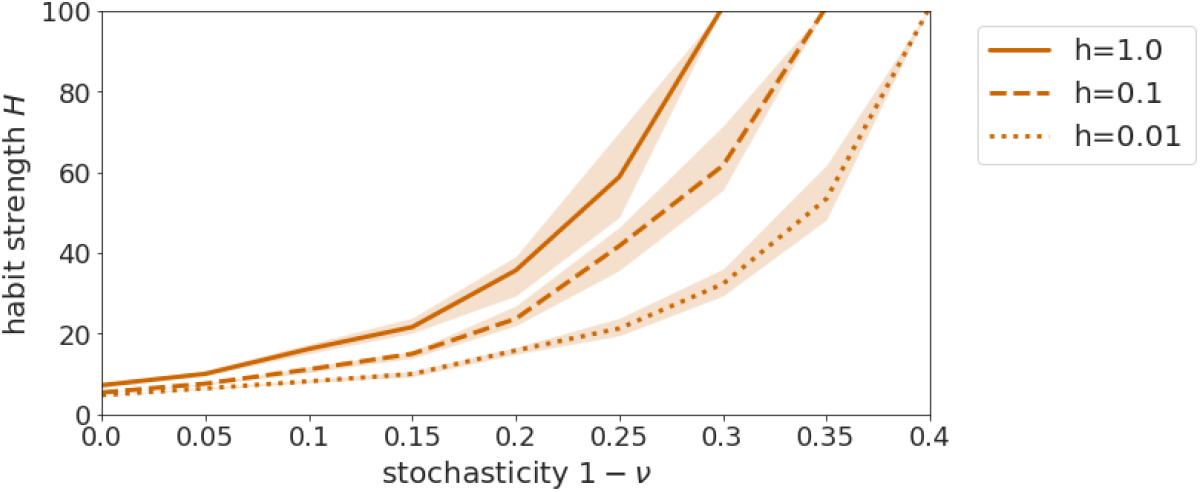
Habit strength as a function of environmental stochasticity 1 – *ν*. The three habit learners (strong, medium, weak) develop stronger habits if the reward scheme is more stochastic, i.e. reward probabilities *ν* are lower. Solid line: strong habit learner with *h* = 1.0; dashed line: medium habit learner with *h* = 0.1; dotted line: weak habit learner with *h* = 0.01. The shaded area surrounding the lines is the confidence interval of 95%. A habit strength of 100 means that the agent does not adapt its behavior within the extinction period of 100 trials.

In the model, this effect is due to two factors: First, as the environment becomes more stochastic, it is harder for an agent to detect the switch contingencies. This delays context inference and thereby action adaptation. Second, the likelihood encoding the goal-directed value is less pronounced in a stochastic environment, as it maps to the decreased probability of achieving a reward for an action. In the model, the agent samples actions from the posterior, which is proportional to the product of the likelihood and the prior. If the likelihood is less pronounced, the habits, as represented by the prior, will automatically gain more weight in the posterior, leading to an increased reliance on habitual behavior in a stochastic environment. Intuitively, this means that a decrease in goal-directed value of actions gives way to a stronger influence of habits. Conversely, habits are also learned more slowly in more stochastic environments because actions are not chosen as consistently because of the decreased goal-directed value. We will come back to the important implications of these findings in the Discussion.

### 3.7. Outcome devaluation

In this section, we show that the prior-based control model can also qualitatively replicate results from outcome devaluation studies, e.g. (Adams, 1982). We modified the habit learning task (Figure 3) by introducing an outcome devaluation in the extinction phase, in addition to the switch in outcome rules. This was done by reducing the prior preference for the reward of lever 1 but not lever 2 in the extinction phase (for details see Appendix).

In general, we find that the outcome devaluation results in a discontinuous jump in the likelihood, as the devalued reward means that action 1 suddenly has no more goal-directed value (data not shown) while action 2 remains useful. Nonetheless, we can apply the same analyses as in Section 3.3 to show the effect of habitual tendency on habit strength under outcome devaluation.

Figure 10 shows, as a function of habitual tendency, (i) the trials numbers in the extinction phase when agents inferred a switch in context and (ii) habit strengths. Independent of habitual tendency (*p* = 0.54, linear regression), agents infer the context switch slightly earlier than in the task without outcome devaluation (median trials 2.4 vs 3.6, *p* < 0.001, two-sample t-test). As before, agents with a low habitual tendency (≤ 0.02) only develop a very weak habit within the training phase of 100 trials (see Figure 4c). When the usefulness of actions now changes due to the devaluation, these agents can instantly, at the beginning of the extinction phase, adapt their behavior to start pressing lever 2. Agents with a higher habitual tendency (≥ 0.1) on the other hand, learn a pronounced habit during training. As a result, these strong habit learners show in the extinction phase after devaluation a delayed action adaptation and thereby a habit strength greater 1 (up to 4). Generally, as before, a higher habitual tendency leads to a greater habit strength (*p* < 0.001, linear regression on the medians).

**Figure 10:**
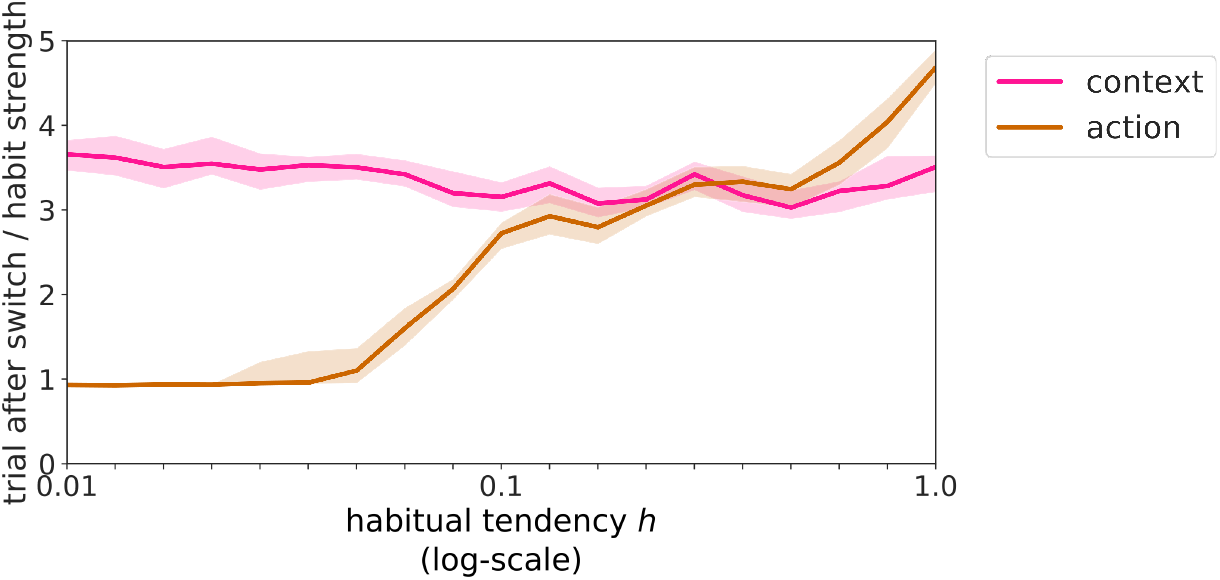
Context inference and action adaptation in the devaluation experiment. Pink line: Time points in the extinction period of when agents infer a switch in context, as a function of the habitual tendency. Brown line: habit strength, as a function of the habitual tendency. The x-axis is logarithmically scaled. This figure is based on the same analysis methods as Figure 5, but here we analyzed the posteriors in an environment with contingency degradation and outcome devaluation.

Clearly, we found a devaluation effect for agents with a habitual tendency *h* ≥ 0.1. Although the habit strengths are fairly low, we found that if we increase the training duration to more extensive training (≥ 500 trials), habit strength increases, so that even weak habit learners show a habit strength greater than 1, and strong habit learners have a habit strength of up to 8 (data not shown).

While these effects are lower than in the contingency degradation experiment, these results show that our model can in principle emulate habitual behavior in both classical experimental designs, contingency degradation and outcome devaluation (Yin and Knowlton, 2006).

### 3.8. Two-step task

Lastly, to show how the model works with sequences of actions, we want to present simulation results for another well known sequential task which has been used to measure habit strength in humans. The two-step task (Daw et al., 2011) rests on the idea that habit learning can be described by model-free reinforcement learning, while goal-directed forward planning can be described by model-based reinforcement learning. Importantly, both systems are weighted to achieve a global control signal for action selection. Concretely, the two-step task is built such that the presumed model-free and model-based control contributions are in conflict under certain conditions. The task (Figure 11) is a sequential decision task with two steps or stages, the first and second level decisions, see (Daw et al., 2011) for details. At the first stage, a subject or agent can choose to visit one of two states, which the subject transitions to probabilistically. There is a common transition with typically a probability of 70% which leads to the state which had been chosen. With a lower probability (30%), a rare transition occurs which leads to the other state instead. At the second stage, a subject or agent can again choose between two options, which will be rewarded with a probability which is subject to a random walk over trials. The availability of the options in the second stage is determined by the state the subject or agent transitioned to from the first stage.

**Figure 11:**
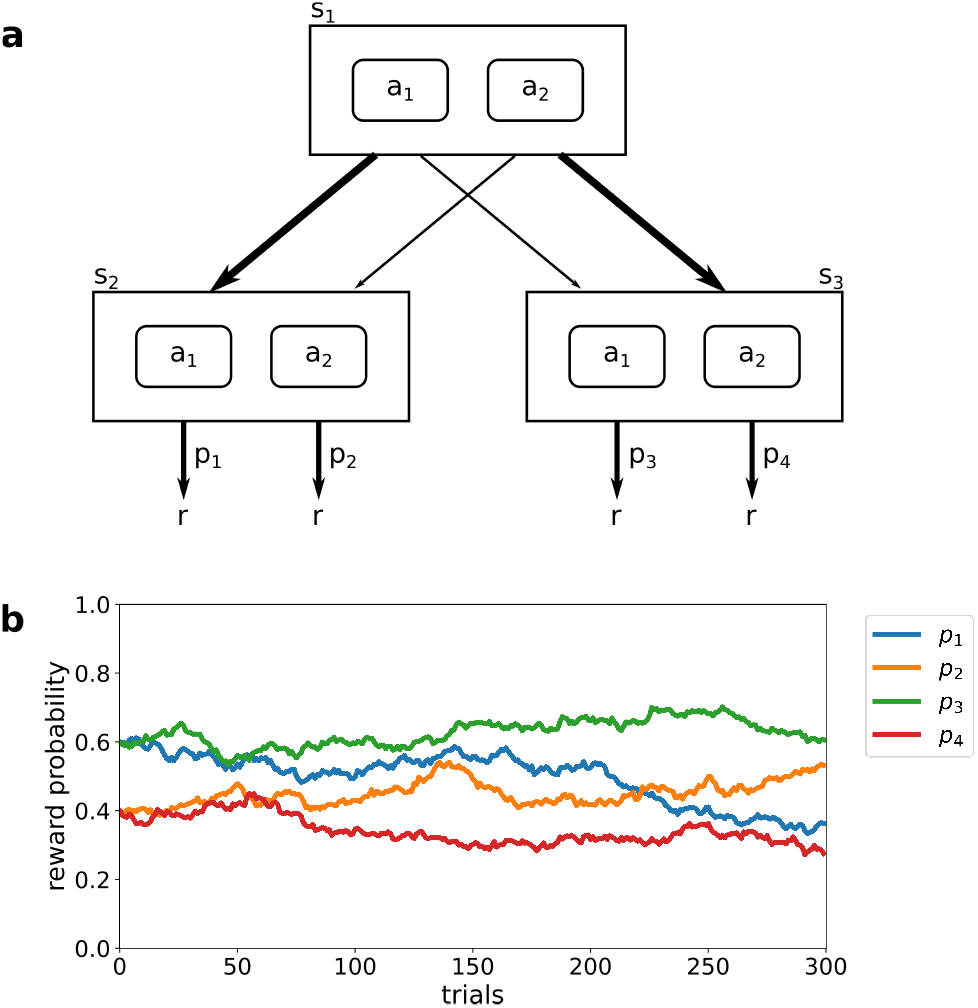
The two-step task. **(a)** Schematic of a trial in the task. The upper rectangle shows the first stage of a trial, where a subject can choose between two actions (*a*_1_ and *a*_2_). For each action chosen, there are two possible transitions to second stage states (*s*_2_ and *s*_3_). Either a common transition occurs with a probability of 70% (thick arrows), or a rare transition with a probability of 30% (thin arrows). The common transition for action *a*_1_ leads to the second state state *s*_2_, and to state *s*_3_ for action *a*_2_, vice versa for the rare transitions. In the second stage of a trial, a subject may again choose between two actions, which may distribute a reward. This makes 4 different actions in the second stage, two of which are only accessible in each state. The four actions have different reward probabilities *p*_1_, *p*_2_, *p*_3_, and *p*_4_ which change over time according to a random walk. **(b)** Random walk of 4 reward probabilities *p*_1_ (blue), *p*_2_ (orange), *p*_3_ (green), and *p*_4_ (red line) as a function of trial number. The figure was adapted from (Daw et al., 2011).

Conflict between the model-free and model-based controller arises after a rare transition: The model-based controller has a representation of the state space and therefore knows that the outcome of the second state choice is more likely to be reached by choosing a different first-stage action. The model-free controller updates action values based on the reward received after having taken this action, independent of whether it is unlikely to get to the same state again using this action. This means that a purely model-based agent is more likely to repeat actions after a common transition leads to a rewarding outcome, or after a rare transition leads to an unrewarding outcome. Equivalently, the purely model-based agent is more likely to switch after a rewarded rare or unrewarded common transition. Conversely, a purely model-free agent is more likely to repeat an action when it has been rewarded, independent of whether a common or rare state transition happened, and is more likely to switch after an unrewarded trial. Experimentally, it is typically found that human participants exhibit a mixture of the two choice patterns, leading to the conclusion that humans employ a mixture of the two strategies, e.g. (Daw et al., 2011).

In order to apply the prior-based control agent to the two-step task, the agent was set up with only one context, as there is no context change in the task. To model the task, we introduced two extensions to the model necessary because of the specific structure of the task: (i) A forgetting factor for the reward as well as the habit learning, which is necessary due to the random walks of the reward probabilities. (ii) A decision temperature in analogy to (Friston et al., 2015) to make the behavioral pattern more pronounced. The values for all three new parameters were set to the values which Daw et al. (2011) inferred from behavioural data, for details of the agent and environment setup see the Appendix.

Figure 12 shows the behavioral patterns of the prior-based control agent. Figure 12**(a)** shows the switch-stay pattern of a purely model-based agent (*h* = 0.0), which corresponds to the typical pattern found from model-based agents, where behavior is repeated more after a rewarded common and unrewarded rare transition. This is not surprising, as model-based agents and the prior-based control agent both rest on a Markov decision process leading to the same goal-directed forward planning. Figure 12**(b)** shows the resulting behavioral pattern of a strong habit learner (*h* = 1.0). As in the original two-step task, behavior is repeated more often after rewarded rare transitions and less often after unrewarded rare transitions. Interestingly and contrary to the classical model-free and model-based agents, our model also predicts a general repetition bias for a strong habit learner in all trial types, as we based habit learning and habitual control strength on repetition of behavior instead of reward and reinforcement. Note that we cannot show a purely habit-based agent here, as such an agent would not fit with the way the prior-based control agent works: An agent requires goal-directed behavior to learn which habits are useful in a specific task environment.

**Figure 12:**
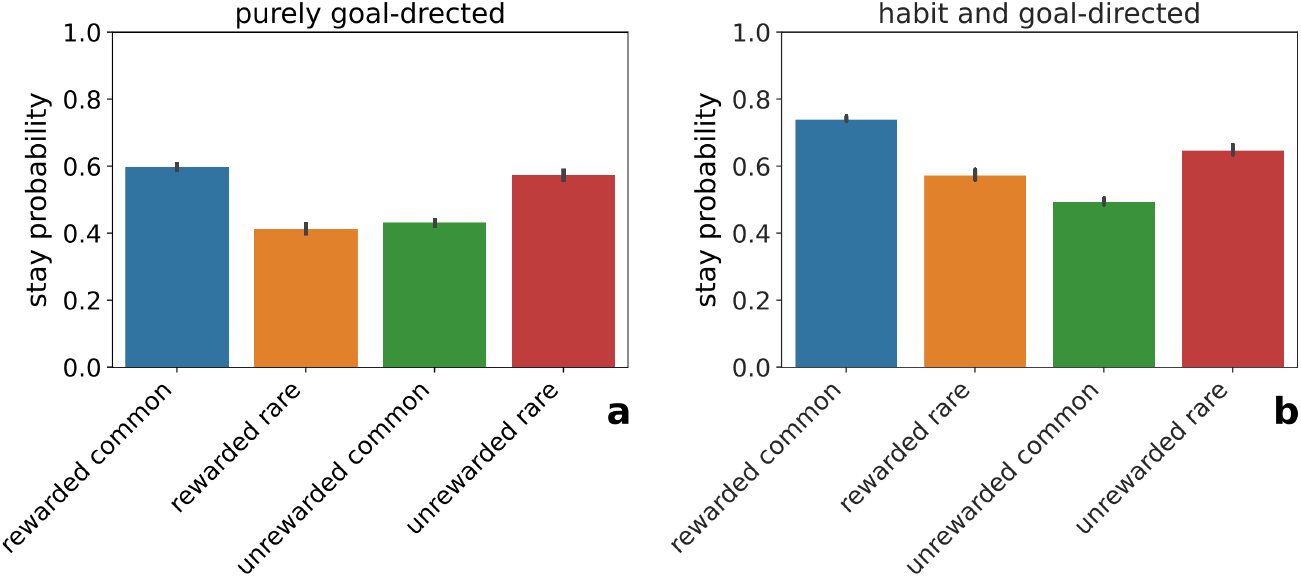
Stay probabilities in the two-step task. **a** Stay probabilities of a purely goal-directed agent, for rewarded common, rewarded rare, unrewarded common, and unrewarded rare transitions (bars from left to right). **b** Stay probabilities of a strong habit learner for different transitions and outcomes. The bar plots are averages of 200 runs.

## 4. Discussion

In this paper, we proposed a hierarchical Bayesian prior-based control model, which may explain how the brain implements habitual learning and balances habitual and goal-directed control. The model rests on four key assumptions about what contributes to habit learning: (i) habits are learned purely based on repetition, (ii) habits can be regarded as chunked action sequences, (iii) control contributions are weighted by their respective uncertainties, and (iv) habits are context-dependent and embedded in a hierarchical model. These assumptions were built into a mathematical model using variational Bayesian methods. Concretely, habitual control contributions were interpreted to stem from a prior over actions or policies, which is learned over time and gives rise to a repetition of previously executed actions or action sequences, implementing assumptions (i) and (ii). Goal-directed control contributions were interpreted to stem from the likelihood, which evaluates a Markov decision process. Actions are then sampled from a posterior, which can be evaluated according to Bayes’ rule and is proportional to the product of the prior times the likelihood, efficiently implementing uncertainty-based weighting (iii) without the need for an additional arbitration unit. This was embedded into a hierarchical model, where an agent can memorize the (habitual) prior, as well as the (goal-directed) action-outcome contingencies (iv) in a context-dependent manner so that an agent can learn and retrieve specific habits and outcome rules for each context it encounters. In short, we interpret experimental evidence for habitual and goal-directed control not as evidence for a dichotomy that competes for action control. Rather, we see action control as a probabilistic inference problem, where three sources of information are integrated: The context, which categorizes the type of situation at hand, the likelihood which evaluates the specifics of this particular instance of situation type, and the prior which is shaped by past experience in this type of situation. Additionally, we used a free (adjustable) parameter to model a trait-like habitual tendency *h*, which determines the learning rate of the prior over policies, and thereby the acquisition speed of the habit.

To show the properties of the model, we introduced a habit learning task with a training and extinction phase, and showed the basic properties of an agent’s information processing employing the model. Using agent-based simulated experiments, we were able to show that our model captures important properties of experimentally established habit learning: insensitivity to both contingency degradation and outcome devaluation, increased habit strength both with extended training duration and with increased environmental stochasticity, and near-instantaneous recovery of habits when exposed to a previously experienced context, as known from ABA renewal experiments. We also found that the habitual tendency interacts with these effects: Agents with higher habitual tendencies exhibit increased habitual contributions to control and habit strength in all of these experimental conditions. Finally, we simulated agent behavior in the two step task where we could replicate the experimentally established behavioral patterns commonly observed in the task.

### 4.1. Interpretation

We identified several causes of variability in habitual control in the agent. First, although habits in the form of a prior over actions are repetition-based, they are formed by the agents goal-directed evaluation while still under goal-directed control. Therefore, they are indirectly influenced by exposure to stochastic stimuli, and their contribution is therefore dynamic and adaptive during a task. In other words, in our model, an agent never stops adapting the prior over policies so that habit strength varies and is context- and experience-dependent. Secondly, we found that behavior is strongly controlled by habits in those situations when goal-directed forward planning cannot determine a clearly best action, so that there is uncertainty on what the best course of action is. This means that habits, when there is conflict between different possible (goal-directed) actions, can be seen as an informed guess to select an action and resolve the conflict rapidly. This uncertainty-weighting of control is in line with previous findings (Daw et al., 2005; Lee et al., 2014). Thirdly, we found that one can emulate a trait-like individual habitual tendency by varying the prior hyper-parameters. The change in hyper-priors is reflected in the change of the individual learning rate during habit acquisition. In turn, the different learning rates are reflected in differences in delayed action adaptation and different habit strengths. Importantly, the habitual tendency parameter not only describes a bias towards one control mode, but inherently describes the time course of the balancing between the control modes. This dynamic balancing is important because it will allow us to not only experimentally test for habits in over-trained individuals, but also in moderately trained individuals, and better understand the time course of habit learning. As such, this parameter is an important free parameter of our model, which can be fitted to data to describe inter-individual differences. For details, we refer the reader to the parameter recovery section in the Supplementary material where we show that the habitual tendency can be inferred from simulated data.

Taken together this leads to an interesting interpretation as to why it may have been advantageous for an organism to evolve to be a strong habit learner with a high habitual tendency. We found that being a strong habit learner supports choosing consistent, reliable behavior, especially in uncertain conditions: When rewarding outcomes were highly stochastic, we found that a habit has a stronger weight on action selection and helps an agent choosing the better option more reliably. Strikingly, precisely this effect has been observed in a recent study, where McKim et al. (2016) found that participants with a history of substance use disorder (SUD) have a heightened ability to execute previously learned stimulus-response associations, in comparison to controls. Assuming that a history of SUD is correlated with a stronger tendency to learn habits, this result directly reflects on our finding of an increased performance for higher habitual tendencies in known contexts. This may mean that being a strong habit learner is advantageous, as long as one’s environment is subdivided into stable phases of already known contexts, separated by infrequent switches. Interestingly, there is evidence for such a mechanism of rapid context-dependent habit retrieval (Bouton and Bolles, 1979; Gershman et al., 2010): Using optogenetics, Smith et al. (2012) observed rapid re-instantiations of a previously learned habit after a context change, where, similar to our simulated experiments, reward contingencies changed. Obviously, this advantage of habits may be even increased, if agents, as is the case in our real-life environment, were able to choose the context they are in or switch to. While we did not implement this active component here, it would most likely lead to agents choosing long stable contexts for which they already learned habits. These scenarios would lead to interesting research about how agents decide to switch contexts to balance exploration and exploitation in their environmental niche.

Even though there are advantages of using habits, it clearly depends on the environment whether habits will be mostly advantageous or disadvantageous. For example, a strong habit learner would be best placed in an environment with rare switches between already learned contexts, see e.g. (Barnes et al., 2005; Gremel and Costa, 2013). Conversely, an environment with frequent changes between contexts dissimilar from previously learned ones would lead to decreased choice performance of a strong habit learner, in comparison to a weak habit learner. Another possibility how the habitual control mechanism may be detrimental to performance is if context inference is for some reason dysfunctional. For example, with suboptimal context inference, one may expect that there is confusion between contexts that are similar in appearance but effectively distinct. We discuss implications of this mechanism in 4.3. We speculate that this confusion may express itself experimentally as an apparent decrease of top-down control by cortical areas (context inference) on the striatum (habitual control), as e.g. found in (Renteria et al., 2018).

Although we argue here that context inference may play a so far underestimated role in habit learning trajectories, context is of course not necessary for modelling habit learning in a task with a single context. This is for example the case for the two-step task used in this work, where an agent exhibits repetition-based habit learning but does not require to switch contexts. Here, another important factor the model comes into play: the learning of habits as chunked action sequences. Dezfouli and Balleine (2013) found that a hierarchical habit learning model using sequences may even fit behavior in the task better than the original model-free model-based habit learning model for which the task was originally devised (Daw et al., 2011). Interestingly, our model predicts a general repetition bias in the two-step task, which sets it apart from the model by Dezfouli and Balleine (2013), and this prediction can be in principle experimentally tested. To leverage the full scope of the proposed model, we hope to, in the future, develop a sequential decision making task for humans which combines all four hypothesized ingredients to habit learning, especially the interpretation of habits as automated action sequences and the context-specificity of habits.

### 4.2. Relation to other habit learning models

In recent years, several approaches to computationally model goal-directed and habitual behavior have been proposed. A prominent interpretation of two distinct habitual and goal-directed systems has been the mapping to model-free and model-based reinforcement learning (Daw et al., 2005, 2011). Here, the model-free system implements an action evaluation based on which actions have been rewarding in the past. The model-based system implements goal-directed forward planning resting on a Markov decision process. Typically, these models have to be run in parallel and require an additional arbitration unit, which evaluates both systems and assigns a weight to each, determining the respective influence on action selection, see e.g. (Lee et al., 2014). This model has often been used in conjunction with the two-step task to asses individual strength of habitual control. However, it seems an open question, whether model-free learning can be indeed mapped to habitual control. For example, Friedel et al. (2014) were able to map model-based reinforcement learning to goal-directed behavior but failed to find such a relation for habitual behavior and model-free reinforcement learning, see also (Wood and Rünger, 2016) for a recent review about the relationship between habitual control and model-free learning.

The prior-based control model differs from this modeling approach in important ways: model-free habit learning proposes that habits arise based on the experienced reward history, while the prior-based control model rests on the assumption that habits arise by repetition. In many experiments, as the contingency degradation experiment used in this work, both assumptions lead to similar experimental predictions. However, our model predicts that habits can arise even in the absence of reward, which model-free learning cannot account for. In addition, model-free habit learning is not hierarchical and does typically not account for context-specific effects. Often, it either lacks contexts completely, or the context is formally implemented as the states at the same level, see e.g. (Palminteri et al., 2015). We propose that behavioral adaptation after a contingency change requires inference of a hidden context, which a typical model-free learning model cannot account for. Instead, the context would have to be known for model-free habit learning to work. Clearly, without context inference, action values would have to be unlearned after a context switch in the environment. In this sense, representing contexts is important for an agent, because behavioural adaptation due to an inferred context change can be very fast, while un- or re-learning is rather slow (Duverne and Koechlin, 2017). Finally, model-based/model-free models usually require an additional arbitration unit, which in turn requires value signals from both controllers. However, if both controllers need to compute values for each action, this approach fails to provide a reasonable explanation how the presumed saving of time and resources can be achieved for habitual control. In contrast, the prior-based control model offers a straight-forward interpretation of how habits can be faster and more resource efficient: When the previously learned prior is close to zero for an action or policy, the posterior is also close to zero. Hence, such a policy will not need to be evaluated by the goal-directed likelihood. In this interpretation, habit learning would equate to learning how to prune the decision tree and confine the policy search space, thereby accelerating the decision process. Future studies will have to investigate whether the prior-based control model can explain reaction-time decreases and resource efficiency of habit learning.

Another intriguing habit learning model resting on the idea of repetition-based habit learning has been introduced by Miller et al. (2018, 2019). The authors proposed to map habitual behavior to a value-free system, which implements a tendency to repeat actions. In this view, the goal-directed system corresponds to a value-based system, which includes model-based as well as model-free reinforcement learning, and both systems are arbitrated using an additional arbitration unit. This proposal is similar to the one in this work as both use repetition-based habit learning, and a Markov decision process for goal-directed control. Indeed, we also deem it reasonable that the brain may employ all behavioral evaluation strategies: value-free, repetition-based learning, as well as model-free reinforcement learning, and goal-directed behavior based on a Markov decision process. The previous work differs from ours in so far as the model requires an additional arbitration unit and needs to evaluate both value-free and value-based signals in parallel. Additionally, and analogously to model-free learning, the model does not include contextual dependencies and context inference. For typical extinction experiments, we would expect both models to make similar predictions. We expect them to differ when an experiment requires context inference and retrieval of previously learned information, like in ABA renewal, which is possible only when habits and action-outcome contingencies are learned in a context-dependent manner.

There have been other Bayesian proposals to habit learning, particularly using active inference. FitzGerald et al. (2014) and Friston et al. (2016) regarded habits in a similar manner to model-free learning, and implemented them as an additional simplified policy. This approach is therefore fundamentally different from and potentially complementary to ours. Nonetheless, we think it possible that the brain uses both value-free as well as model-free learning processes, so that it would not be unreasonable to assume that both contribute to action selection. Maisto et al. (2019) regarded habits as cached values of the likelihood calculated in previous trials of the experiment. This means that the likelihood was only calculated when first encountering a new context, and is then kept stable and cached as long as the context does not change. These proposals of a Bayesian treatment of habit learning are different from (and possibly complementary to) our approach, as we view habits as a prior over actions or policies, and not related to the likelihood (which in our model represents goal-directed control). Under extensive training regimes, both approaches might lead to similar results. However, under limited training, when both, goal-directed and habitual control influence behavioral control, our approach may lead to more plausible behavior in this regime because of the balancing of the two contributions.

Furthermore, there are other proposals to view reinforcement, contingency degradation, and outcome devaluation experiments as a context inference and rule learning problem (Palminteri et al., 2015; Gershman et al., 2010; Wilson et al., 2014). These studies view task states as latent variables or contexts, which need to be inferred, while reward generation rules from these states are learned, which essentially translates to a non-hierarchical, partially observable Markov decision process. What sets our proposal apart, is that we view the context as a latent variable on a higher level of a hierarchical model, which modulates how rewards are generated from the same states. This allows us to describe not only actions but sequences of actions which enables an agent to navigate a state space, where the rules might change even within the same environment. We can thereby incorporate the assumption that habits are based on chunked action sequences which allows us to map our model to interesting neurophysiological findings which we discuss below. These ideas also align with proposals such as event coding (Hommel et al., 2001) and event segmentation theory (Zacks et al., 2007), which posit that behavior is segmented into events or episodes. Based on these proposals, Butz et al. (2019) put forward an interesting context and contingency learning model which implements ideas similar to our goal-directed evaluation in a neural network model.

### 4.3. Implications

A topic highly related to the interaction between goal-directed and habitual control is substance use disorder (SUD): previous accounts have described for SUD a shift from goal-directed to habitual and compulsive use (Everitt and Robbins, 2005). One interesting aspect of the proposed model is that context inference may have a much more central role in this shift to habitual control. Here we discuss briefly the potential implications of a dysfunctional context inference mechanism for addictive behavior. Our proposed model combines repetition-based habit learning with context inference, where delayed context inference leads to prolonged repetition of behavior even when previous actions are not rewarded. We speculate that both these aspects could play a pivotal role in substance use trajectories and may help to explain why it can be hard to stop substance use even in the face of adverse consequences. Contrary to the typical extinction experiments, real life action-outcome contingencies associated with use do not switch suddenly, instead outcomes go slowly from being initially rewarding to bearing negative consequences. Such a slow transition may make it hard to update beliefs about the context, as there is no definite switch, which would prompt behavioral adaptation and the stopping of use. Additionally, the habitual control contributions in our model are based on repetition alone, which leads to an action being repeated despite negative outcomes which can only slowly be unlearned, unless a new habit is learned in a new context. These dynamics fit well to studies from social psychology, e.g. (Lally et al., 2010; Danner et al., 2008; Neal et al., 2012; Verplanken and Roy, 2016), which show that every-day habits are learned more quickly in a consistent context, but can be unlearned more easily after a context change.

Another interesting and experimentally relevant example of biased context inference may be the established phenomena of Pavlovian to Instrumental Transfer (PIT), (Garbusow et al., 2014; Talmi et al., 2008), where participants are biased towards a previously encountered context by cues of that context. Although our model does currently not include explicit cues, such a generalization can be implemented in future version and would allow the application to PIT tasks. In terms of the prior-based control model, Pavlovian learning in specific PIT would imply learning reward-context associations, while instrumental learning corresponds to learning of outcome contingencies. We speculate that these two parts of a PIT experiment could be represented in the brain as two contexts, where specific rules for each of these environments would be learned. When participants then re-encounter the conditioned stimulus from the Pavlovian learning during instrumental behavior, their context inference becomes biased towards this previously experienced context, introducing uncertainty on beliefs about contexts. As a result, both representations would be loaded at the same time, which the brain may then try to map onto each other, leading to increased approach behavior when the previously conditioned stimulus is present.

In summary, the proposed modelling approach provides the novel perspective that habitual control relies on learned context-specific priors of policies. The resulting model provides for a simple way to balance action control between habitual and goal-directed control. As we have discussed, experimental findings seem to support this perspective of a separation into prior and posterior over policies. We anticipate that the present computational modelling approach may support novel directions of research aimed at the central role of context inference as a means to reduce the number of policies that have to be evaluated and implement fast action control relying on the interplay between the prior and posterior over policies.

## 5. Acknowledgments

We thank Ann-Kathrin Stock for valuable comments and suggestions.

## 6. Funding acknowledgments

Funded by the German Research Foundation (DFG, Deutsche Forschungs-gemeinschaft), SFB 940/2, projects A9 and Z2, and TRR 265, project B09.

## AppendixA. Appendix

### AppendixA.1. Python code

As described in the main text, the update equations, artificial agents, and experimental environments were implemented in python. The code was made publicly available in the following github repository: https://github.com/SSchwoebel/BalancingControl

We invite any interested reader to download, use, or modify the code, and to contact us should any questions arise.

### AppendixA.2. Derivations of the update equations

The variational free energy functional is defined as the Kullback-Leibler divergence between the approximate posterior 14 and the joint probability distribution of the generative model 6. Hence, we can write the variational free energy as

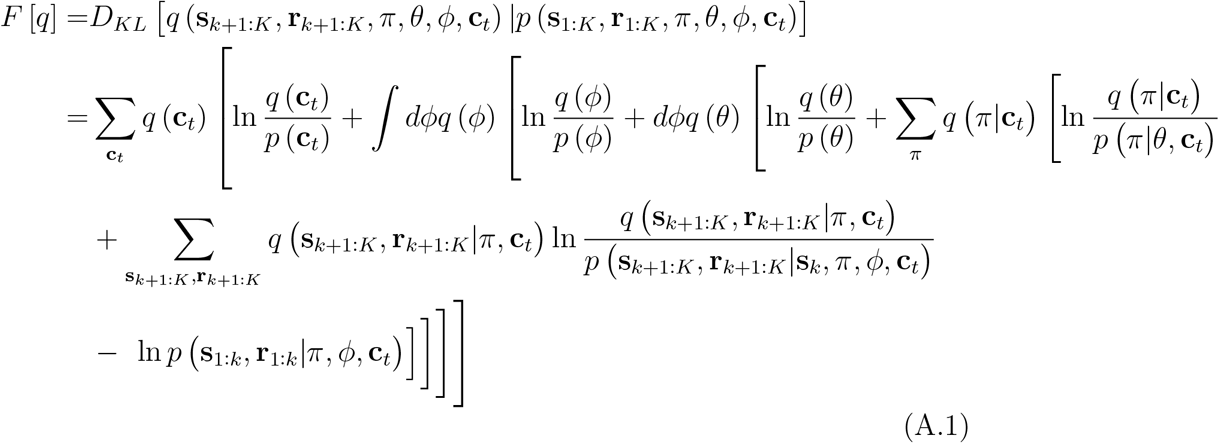

where for clarity we omitted the parametric dependence of each distribution. The approximate posterior is than obtained as the minimum of the free energy, defining the upper bound on surprise (negative marginal log-likelihood).

We first write down the update equations for the beliefs over future states and rewards within an episode, using the belief propagation message passing update rules (Pearl, 2014; Yedidia et al., 2003). For details on the derivation steps see our previous work (Schwöbel et al., 2018) in which we investigated the Bethe approximation for a Bayesian treatment of a partially observable Markov decision process. The results shown here are an adaptation for fully observable states, which is just a special case.

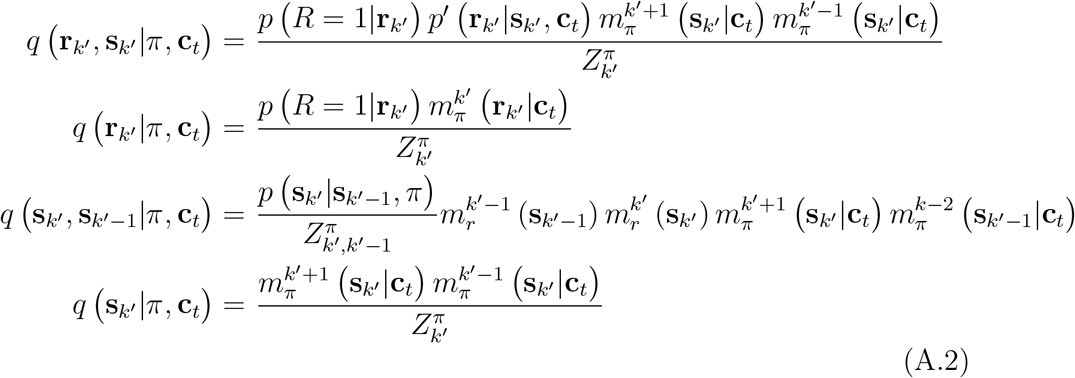

using the messages

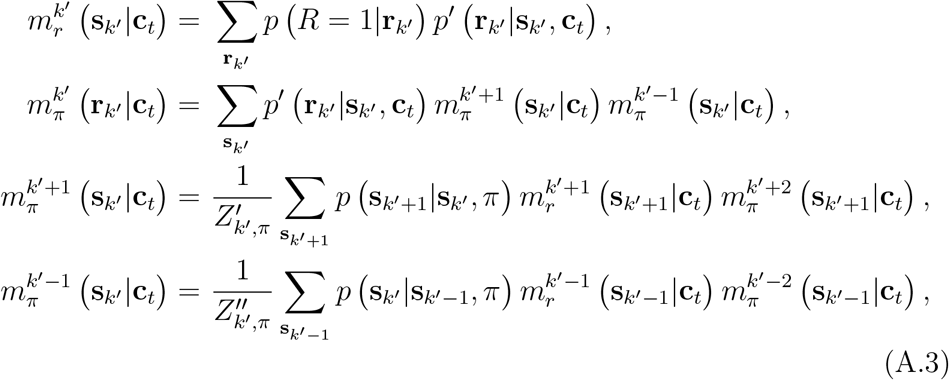

where

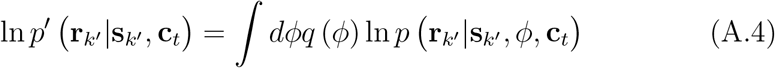

the free energy mandated that we average out the dependency on *ϕ*.

The posterior beliefs over policies given some context are calculated as

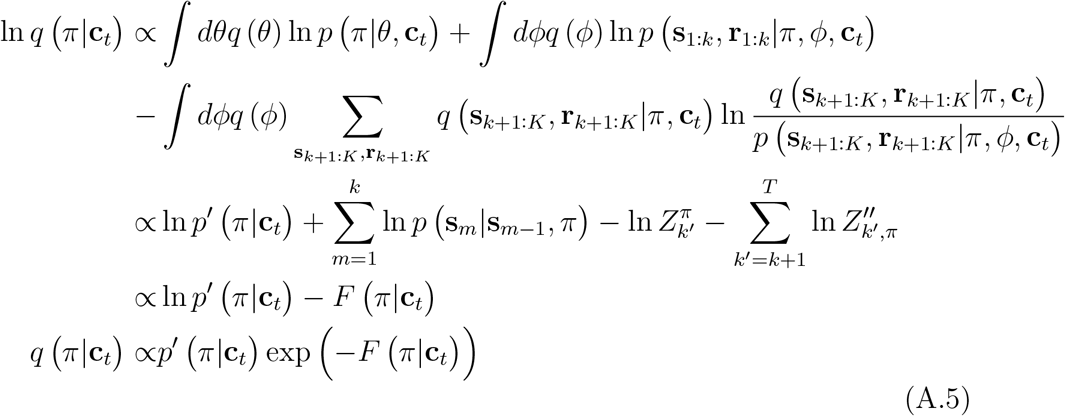

where *p′*(*π*|**c**_*t*_) is the marginalized prior over policies, and *F*(*π*|**c**_*t*_) is the policy-specific free energy in a given context (see (Schwöbel et al., 2018)).

The posterior over the parameters *θ* of the prior over policies can be derived as

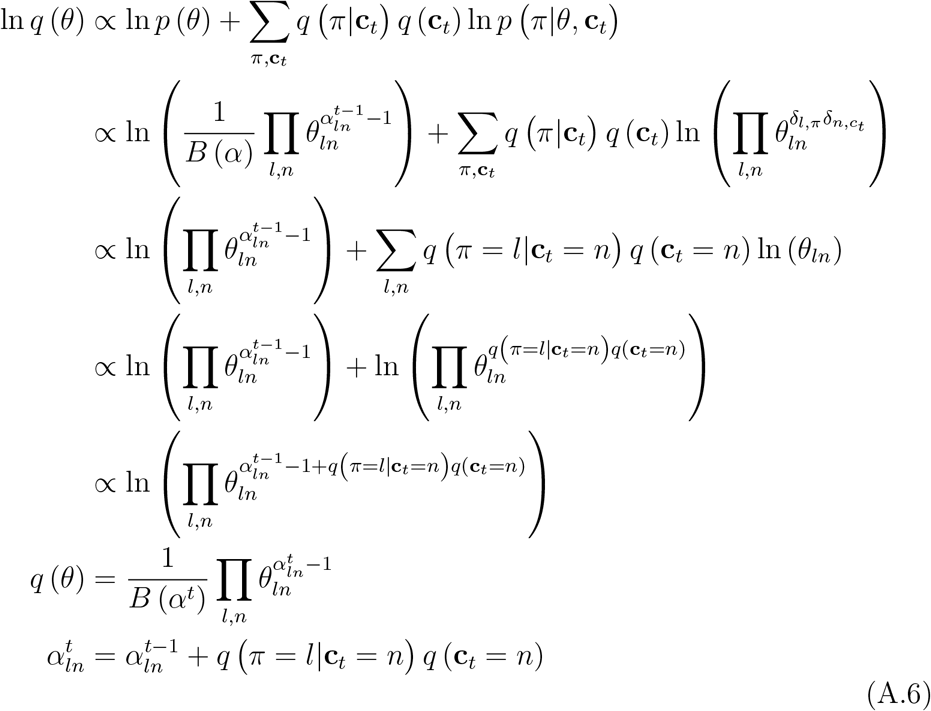

and is itself again a Dirichlet distribution with updated pseudo counts *α^t^*. These are updated by adding the posterior over policies times the posterior over context. At the end of an episode, the pseudo count will be increased by 1 for the policy which has been followed in the context the agent inferred to be in.

The posterior over the parameters *ϕ* of the outcome rules can be derived as

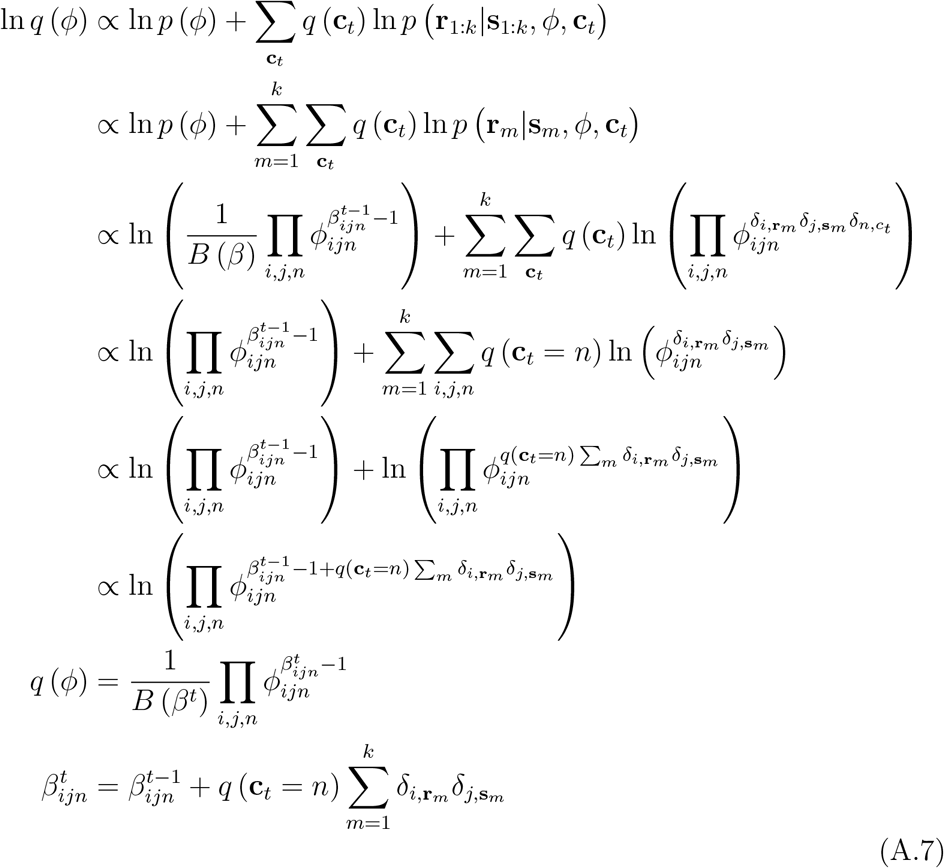

Lastly, we want to derive the posterior over contexts

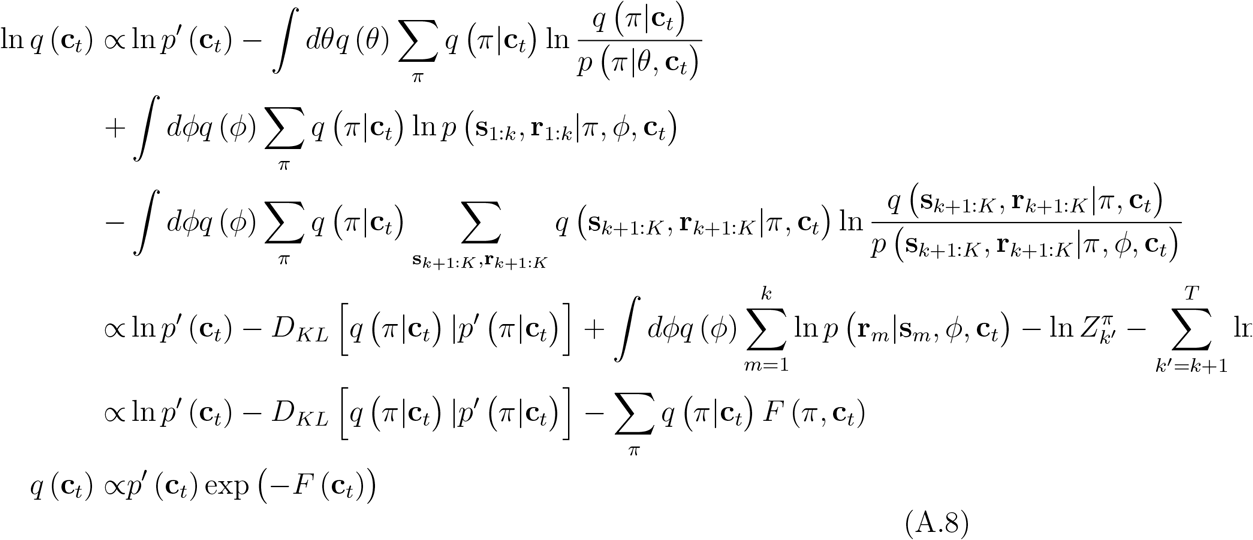

with context-specific free energy *F*(**c**_*t*_). Note, that we set

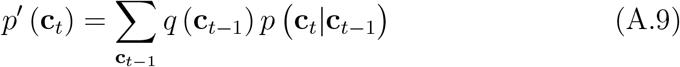

As most of the posteriors described here are interdependent on each other, one has to iterate over their updates until convergence. Practically, we only used one iteration step: We used the priors over *θ, ϕ* and **c**_*t*_ to calculate the posterior over policies. Then we calculated the posteriors over *θ* and *ϕ*, which were then used to calculate the posterior over contexts. We evaluated if this procedure is equivalent to a full iteration until convergence and found that the resulting posteriors only differed by less than 1% of their values.

### AppendixA.3. Agent and extinction task setup

The generative process of the habit learning task (Section 3.2) was set up as follows:

- An episode has length *K* = 2.
- There are 200 episodes so that *t* ∈ [1,200]
- There are *n_s_* = 3 states 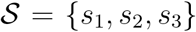, where *s*_1_ is the state where lever 1 distributes a reward, *s*_2_ is the state where lever 2 distributes a reward, and state *s*_3_ is the starting state in front of the two levers.
- There are *n_r_* = 3 rewards 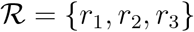, where *r*_1_ is the reward payed out by lever 1, *r*_2_ is the reward payed out by lever 2, and *r*_3_ is the no-reward.
- There are *n_a_* = 2 actions 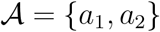, where *a*_1_ leads to state *s*_1_, and *a*_2_ leads to state *s*_2_ from any starting state.
- There are *n_c_* = 2 contexts 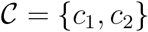 which amount to lever 1 or lever 2 being the better arm, respectively.
- The state transitions are set up to be deterministic:

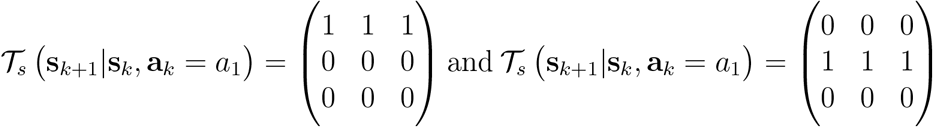

so that *a*_1_ leads to state *s*_1_ from any starting state, and *a*_2_ to *s*_2_, while *s*_3_ can not be reached.
- The reward generation rules are as depicted in Figure 3b. Mathematically, the reward generation in the training phase as 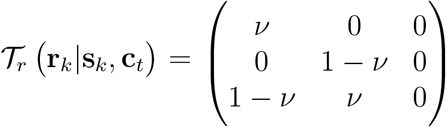 for *t* ∈ [1, *d_training_*], where *ν* is the probability of lever 1 distributing a reward. In the extinction phase, the reward probabilities switch, so that 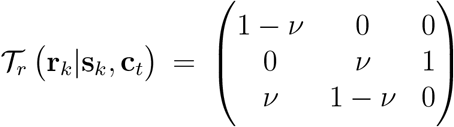 for *t* ∈ [*d_training_* + 1; *d_training_* + 100]
- The context transitions are deterministic and happen after the training, so that 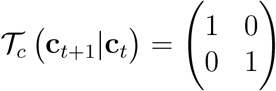 for *t* ∈ {1, …, *d*_training_, *d*_training_ + 2, …, *d*_training_ + 100} and 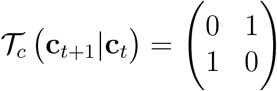 for *t* = *d*_training_ + 1.

In each episode *t*, the agent starts at *k* =1 in the state *s*_3_ in front of the levers.

The agent’s generative model is set up to reflect the generative process, or learn the respective quantities:

- The agent knows it starts in state *s*_3_ in each episode, so we set the prior of the starting state as *p* (**s**_1_|**s**_0_, *π*) = *p* (**s**_1_) = (0, 0,1)*^T^*
- As we set *K* = 2, policies and actions map one to one, so that *len* (*π*) = 1 and *n_π_* = 2. This means, *π*_1_ = *a*_1_ and *π*_2_ = *a*_2_
- We assume the agent knows the state transitions instead of learning those, so that 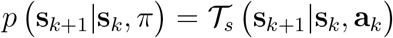
- The pseudo counts 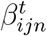 which are used to parameterize the outcome rules for reward *i* and state *j* in context *n*, are initialized as 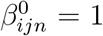 for all *i*, *j*, *n*
- The pseudo counts 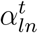 which parameterize the prior over actions for policy *l* in context *n* are initialized as 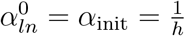 using the habitual tendency *h* and are initialized the same for all *l*, *n*.
- We set the agent’s representation of context transitions, i.e. temporal stability as 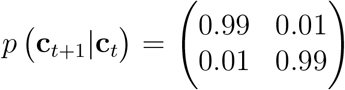 for *t* = *d*_training_ + 1. Here, the agent assumes that both contexts are equally stable and change once in 100 trials.
- Finally, we set the agents preference for outcomes as *p* (*R* = 1|**r**_*k*′_) = (0.495, 0.495, 0.01)*^T^*, so that the agent prefers the rewards of levers 1 and 2 equally, but dislikes the no-reward *r*_3_. In the contingency degradation tasks, these values are kept constant. In the outcome devaluation task (Section 3.7), the preference for outcomes was reset in the extinction phase as *p* (*R* = 1|**r**_*k*′_) = (0.01942, 0.96117, 0.01942)*^T^*, which effectively devalues the reward for lever 1 and keeps the ratio of desirability between reward and no reward unchanged.

### AppendixA.4. Two-step task setup and update equation modifications

The generative process of the two-step task was set up as follows:

- An episode has length *K* = 3.
- There are 300 episodes so that *t* ∈ [1, 200]
- There are *n_s_* = 7 states 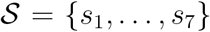. *s*_1_ is the starting state, *s*_2_ and *s*_3_ are the first stage states, *s*_4_, and *s*_6_ are the second stage states corresponding to the choice options in state *s*_1_, and *s*_5_, and *s*_7_ are the second stage states corresponding to the choice options in state *s*_7_.
- There are *n_r_* = 2 rewards 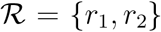, which correspond to the no-reward and reward, respectively.
- There are *n_a_* = 2 actions 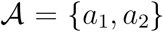, which correspond to the two choice options at each state.
- There is *n_c_* =1 context 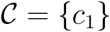.
- The state transitions are probabilistic:

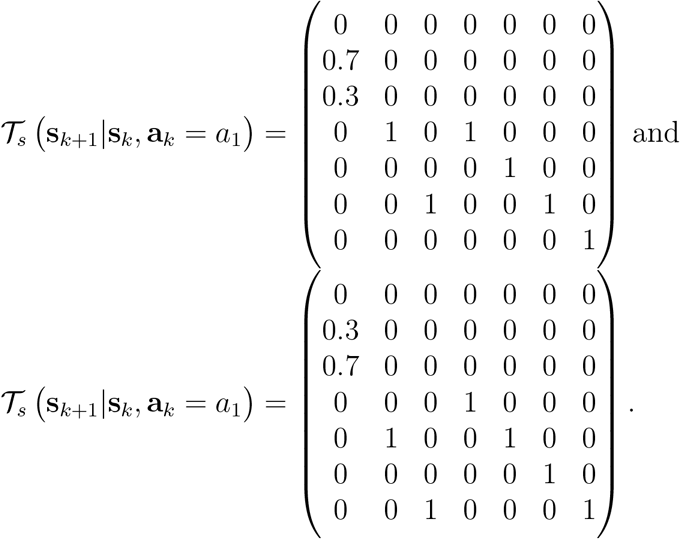
- The reward generation rules correspond to a random walk and are as depicted in Figure 11b.

The agent’s update rules were modified in 3 ways: Firstly, we introduced a decision temperature *γ* in analogy to (Friston et al., 2015). The modified posterior over policies is

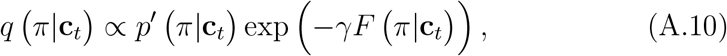

where we set *γ* = 4, which corresponds to the mean decision temperature Daw et al. (2011) found for participants in their original paper.

Secondly, we introduced a forgetting factor λ_1_ in the learning of the concentration parameters *α* of the prior over policies. The modified update equation reads

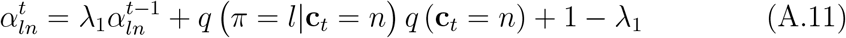

where we set λ_1_ = 0.7, again in accordance with the parameters Daw et al. (2011) found in their participants. We refer the interested reader to (Moens and Zénon, 2019) and (Lu et al., 2019) for work on how to introduce forgetting factors into Bayesian updating.

Lastly, we introduced a forgetting factor λ_2_ to the learning of the reward contingencies. The modified update equation is

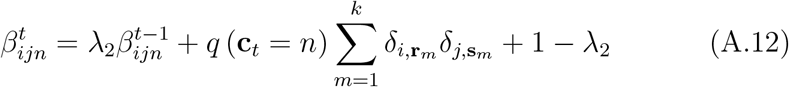

where we set λ_2_ = 0.4 in accordance with (Daw et al., 2011)

