## Supplementary material for "Balancing control: a Bayesian interpretation of habitual and goal-directed behavior"

**Simulations with 3 contexts.**

Here, to show that the model is not constrained to just two contexts, as a proof of principle we show how the agent fares with three contexts. In order to focus on context inference, we chose the weak habit learner with a habitual tendency $h = 0.01$ for the following simulations. A straight-forward choice for subjecting an agent to $m \geq 3$ contexts is to generalize the experiment to a multi-armed bandit problem. Figure 1 shows such a generalization for $m = 3$ arms. The task was set up similar as in the main text, so that each arm has a phase of 100 trials where it has the highest reward probability of $\rho = 0.9$ (Figure 1(**a**)). The agent infers contexts (Figure 1(**b,c**)) and actions (Figure 1(**d,e**)) well and learns to associate each context with the respective action-outcome contingencies. Note that the agent has no prior information about what levers or actions are associated with which context. Instead, the agent infers to be in a new context once the outcomes in the environment become sufficiently surprising, given its previous knowledge about the environment. This lack of prior information is illustrated in Figure 1(**c,e**), which shows that ordinal numbers of contexts are not matched to the ordinal numbers of best levers in that context.

To put the model further to the test, we also introduced an environment with 2 levers but $m = 3$ contexts (Figure 2(**a**)). The first two contexts, which run over 200 trials in this experiment, are the same as in the main text: The first 100 trials, lever 1 is optimal with a reward probability of $\rho = 0.9$, the second 100 trials lever 2 is optimal with the same reward probability. Additionally, there is now a third phase in the last 100 trials, where both levers distribute rewards randomly. The third phase should be qualitatively different enough for an agent to infer that this is a third context. And indeed, the context inference (Figure 2(**b,c**)) shows that the agent successfully infers that the last phase is a novel third context. Note that in all simulations shown in this section we fix the maximal number of possible contexts to $n_c = 3$.

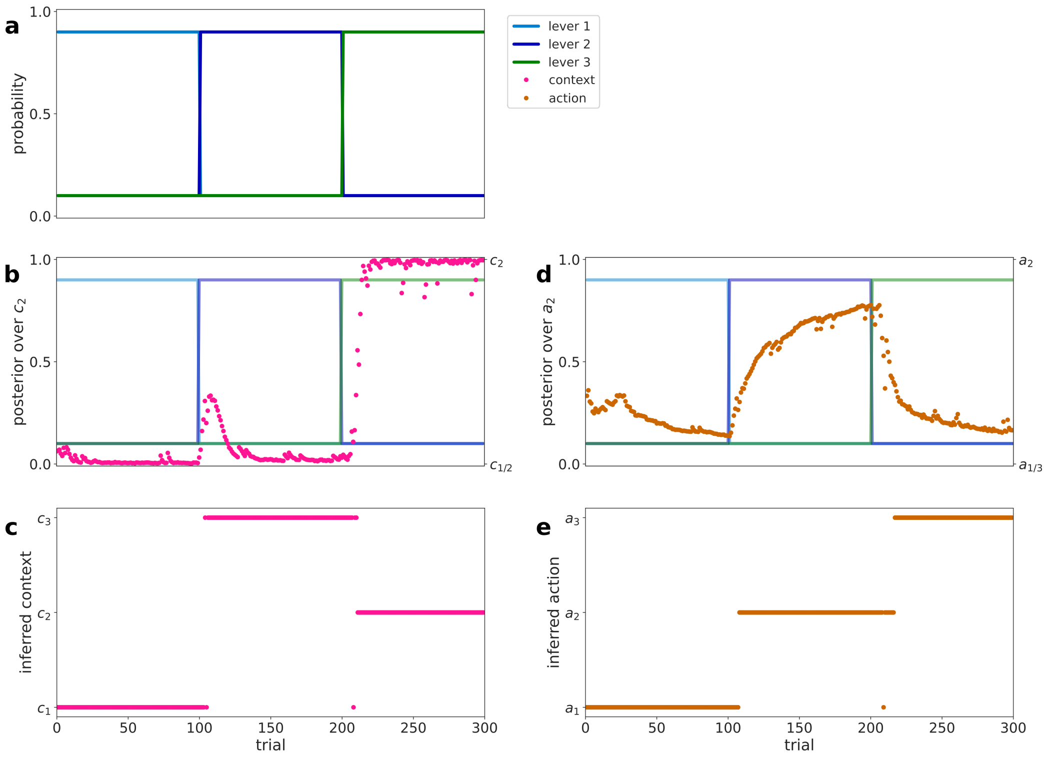


Figure 1: Task with 3 levers

**(a)** Modified three-contexts task: The agent has three options (levers) where we denote the time course of reward probabilities associated with each lever with light blue, dark blue, and green lines, respectively. Only one of the levers is associated with the highest reward probability of $\rho = 0.9$ at each time. This configuration is fixed for duration of 100 trials, after which the optimal lever changes. **(b)** The inferred posterior probability that the agent is in context 2 (pink dots). **(c)** The maximum a posteriori estimate over contexts (pink dots), i.e. which context the agent inferred to most likely be in. Note that the context number and lever number are not necessarily matched. However, each context is associated with one best lever. The right column (**(d,e)**) shows the agent's action inference. **(d)** The posterior over action 2 (brown dots), of which the agent learns over the time course of the experiment that it is the optimal action when lever 2 has the highest reward probability. **(e)** The maximum a posteriori estimate over actions (brown dots), i.e. which actions the agent deemed optimal at any trial. The agent successfully infers that actions 1,2,3 are best when levers 1,2,3 have the highest reward probability, respectively. It also learned to connect action 2 with context 3 and action 3 with context 3.

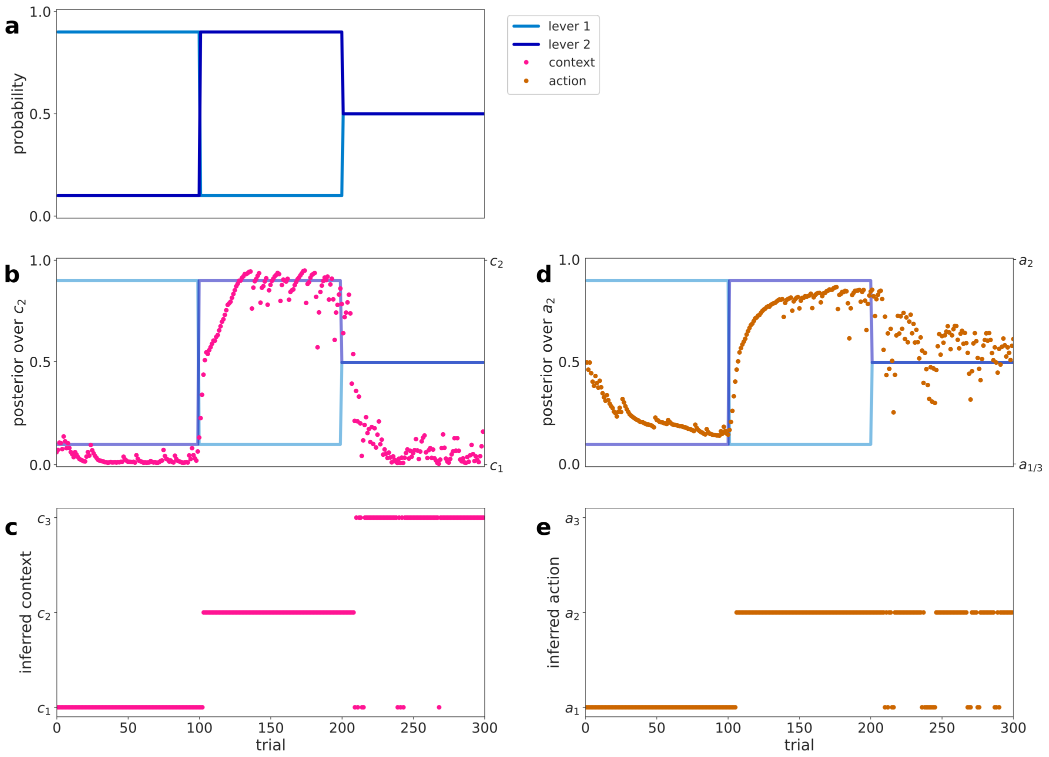


Figure 2: Task with 2 levers and 3 contexts

**(a)** The modified task structure: Starting from trial 200, both levers distribute rewards randomly with the respective probabilities as 0.5, in contrast to the first two contexts. This constitutes the third context, as this configuration is qualitatively different from the others. **(b)** shows the posterior over context 2, and **(c)** shows the maximum of the posterior over contexts (pink dots), i.e. which context the agent inferred to most likely be in. The agent successfully classified all three phases as thee different contexts. The right column shows the agent's action inference. **(d)** shows the posterior over action 2 (brown dots), of which the agent learns over the time course of the experiment that it is the optimal action when lever 2 has the highest reward probability. The posterior over actions goes towards 0.5 in the third phase of the experiment, where both actions distribute reward with equal probability. **(e)** shows the maximum of the posterior over actions (brown dots), i.e. which actions the agent deemed optimal at any trial. The agent learns that action 1 is best in the first phase of the experiment, and action 2 is best in the second phase. In the third phase, as the posterior over actions goes towards 0.5, the agent randomly infers one action to be better than the other.

### Parameter recovery

In this section we demonstrate the inference of the trait-like habitual tendency parameter from simulated data. Due to learning effects, and low sensitivity to initial conditions, it is not always possible to recover the underlying habitual tendency of a single simulated participant. Instead, we will show here that it is possible to recover the habitual tendency $h$ on a group level, and tease apart behavior that was generated by agents with low ($h = 0.01$), medium ($h = 0.1$), and high ($h = 1.0$) habitual tendencies.

To implement this, we set up five groups consisting of $N = 20$ simulated participants (agents), where we fix habitual tendencies of all agents within a group to the same, group specific value. The group specific habitual tendencies correspond to $0.01, 0.033, 0.1, 0.33$, and $1.0$. Each of these groups performs the extinction task in two exemplary environments: one with a low reward probability $\rho = 0.75$ and moderate training duration $d_{\text{train}} = 100$, and one with a larger reward probability $\rho = 0.9$ and a long training duration $d_{\text{train}} = 100$. Each agent generates $T = 200$ responses in the two experimental conditions.

The group level posterior estimates of habitual tendencies for each group and each environment are shown in Figure 3. These two environments were chosen based on the highest divergence of the resulting habit strength (see Figures 6 and 9 in the main text). In these two environments, the group-level habitual tendency could be recovered reasonably well, specifically, it is possible to differentiate between groups of high, medium and low habitual tendency. This property of the model will be important when this model is applied to future experimental studies. For an environment with high reward probability and moderate training duration, as used in the main text in Figure 3, the group level habitual tendency could not be recovered (not shown), and inference usually yielded low habitual tendencies, even for groups with high habitual tendencies.

In the following we show how we set up the parameter recovery and inference process formally. Each group in an environment yields a data set $D = \{D_1, \dots, D_N\}$ consisting of choice-outcome data tuple $D_i = (\mathbf{a}^i_{1:T}, \mathbf{r}^i_{1:T})$ for each simulated agent $i$ and for $T$ trials. We define the response likelihood function as

$$p\left(D_i|h\right) = p\left(\mathbf{a}^i_{1:T}|\mathbf{r}^i_{1:T}, h\right) = \prod_{t=1}^{T} q\left(\mathbf{a}^i_t|h\right) \tag{1}$$

where $q\left(\mathbf{a}^i_t|h\right)$ denotes posterior action probability of the $i$th agent on trial

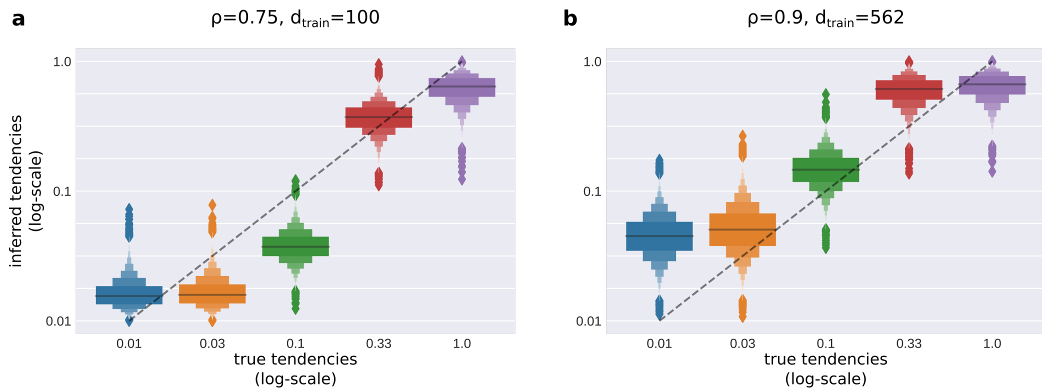


Figure 3: Parameter recovery

**(a)** Inferred habitual tendencies for an extinction task with a reward probability of $\rho = 0.75$ and a training duration of $d_{\text{train}} = 100$ trials. The true habitual tendency used for the generation of simulated behavioral data is shown on the x-axis, and the y axis shows the inferred habitual tendency. The bars for each true habitual tendency indicate the inferred distribution. The dashed line indicates the true habitual tendency for each group. In inference, the true habitual tendency can be inferred, and data generated with low, medium, and high habitual tendency can be differentiated. **(b)** Inferred habitual tendencies in a task with higher reward probability ($\rho = 0.9$) and long training duration ($d_{\text{train}} = 562$). The plot is as in **(a)**. Under this longer training duration, the underlying habitual tendency of a group can be inferred well.

$t$ after observing a sequence of rewards $\mathbf{r}^i_{1:t}$ and performing a sequence of choices $\mathbf{a}^i_{1:t-1}$, for a fixed habitual tendency $h$. To facilitate inference, we constrain habitual tendencies to $M = 9$ categories $c \in \{0, \ldots, 8\}$; hence $h_c \in \{0.01, 0.018, 0.032, 0.056, 0.1, 0.18, 0.32, 0.56, 1.0\}$, and $h_c = 10^{-\frac{c}{4}}$. Here, we chose the categories so that they are equidistant in log-space. The hierarchical generative model of behavioural data is defined as

$$\begin{aligned}
\alpha_j &\sim \Gamma\left(\alpha = 1, \beta = 0.5\right), \ \text{ for } j \in \{1, \ldots, M\} \\
\vec{p} &\sim Dir\left(\vec{\alpha}\right) \\
c_i &\sim Cat\left(\vec{p}\right) \\
D_i &\sim P(D_i|c_i)
\end{aligned} \tag{2}$$

where, we assume a random-effect hierarchical model, with hyper-priors set to Dirichlet and Gamma distributions.

Effectively, the main interest here is to infer the posterior of the probability vector $\vec{p}$, which defines the group level distribution of habitual tendencies. Hence, we marginalize out the categories and obtain the following marginal likelihood

$$p\left(D_i|\vec{p}\right) = \sum_c p(D_i|c)p_c \tag{3}$$

which we can utilise to obtain the posterior distribution over hyper-priors (group level parameters) as

$$p\left(\vec{p}, \vec{\alpha}|D\right) \propto p\left(\vec{p}|\vec{\alpha}\right) \prod_j p(\alpha_j) \prod_i p\left(D_i|\vec{p}\right) \tag{4}$$

Finally, given a sample from the marginal posterior distribution $\vec{p}_n \sim p\left(\vec{p}|D\right)$ we obtain a corresponding group level sample of habitual tendency as

$$h_n = \sum_c f\left(c\right) p^n_c. \tag{5}$$

We implemented the generative model (Eq. 4) in PyMC3 probabilistic programming package (Salvatier et al., 2016), where we use a NUTS sampler for generating approximate samples from the posterior. For details of the implementation, we refer the reader to our github repository (https://github.com/SSchwoebel/BalancingControl).
